# Pro-Resolving Mediator Profiles And 5-Lipoxygenase Activity In Cerebrospinal Fluid Correlate with Disease Severity and Outcome in Adults with Tuberculous Meningitis

**DOI:** 10.1101/608901

**Authors:** Romain A. Colas, Le Thanh Hoang Nhat, Nguyen Thuy Thuong Thuong, Esteban Alberto Gomez Cifuentes, Lucy Ly, Hai Hoang Thanh, Nguyen Hoan Phu, Nguyen Thi Hoang Mai, Guy E. Thwaites, Jesmond Dalli

## Abstract

Tuberculous meningitis (TBM) is the most lethal form of tuberculosis infection, characterized by a dysregulated immune response that frequently leads to neurological injury and death despite the best available treatment. The mechanisms driving the inflammatory response in TBM are not well understood. To gain insights into these mechanisms we used a lipid mediator profiling approach to investigate the regulation of a novel group of host protective mediators, termed specialized pro-resolving mediators (SPM), in the cerebrospinal fluid (CSF) of adults with TBM enrolled into a randomised placebo-controlled trial of adjunctive aspirin treatment. We found distinct lipid mediator profiles with increasing disease severity, changes that were linked with an upregulation of inflammatory eicosanoids in patients with severe TBM and a decrease in the production of a number of 5-lipoxygenase (ALOX5)-derived SPM. CSF pro-resolving mediator concentrations were also associated with 80-day survival. In survivors, we found a significant increase in pro-resolving mediator concentrations, including the ALOX5-derived resolvin (Rv)T2, RvT4 and 15-epi-Lipoxin (LX)B_4_, compared to those who died. Aspirin administration increased the ratio of pro-resolving to pro-inflammatory mediators decreasing the concentrations of the prothrombic mediator TxA_2_, changes that were linked with early reductions in brain infarcts and deaths. Together, these findings identify a CSF SPM signature that is associated with disease severity and 80-day mortality in TBM. Furthermore, the therapeutic manipulation of the ratio between pro-resolving mediators and pro-inflammatory/thrombogenic mediators in the CSF, by aspirin for example, offers a novel treatment strategy to reduce the morbidity and mortality caused by TBM.

**Authors Summary:** Infections of the brain and the meninges by *Mycobacterium tuberculosis* (*M. tb*) lead to severe inflammation and are associated with poor outcomes. The mechanisms leading to this disease remain poorly defined. Herein, we investigated how *M. tb* infection regulates the concentrations of specialized pro-resolving mediators that are central in controlling the body’s ability to clear infections. In these investigations, we found that disease survival was linked with increased concentrations of a number of these protective molecules including resolvins and lipoxins. Treatment of *M. tb*-infected patients with aspirin decreased the production of the immunosuppressive and thrombogenic mediator thromboxane A_2_ improving the balance between protective and inflammatory molecules. Of note, these changes were linked with reduced disease severity and improved survival. Therefore, the present findings suggest a previously unappreciated role for pro-resolving mediators in TBM pathogenesis.

## Introduction

*Mycobacterium tuberculosis* (*M. tb*) is responsible for more deaths globally than any other infectious disease. When it infects the brain and meninges to cause tuberculous meningitis (TBM), which represents 1-5% of all forms of tuberculosis, it either kills or severely disables around a half of all sufferers despite the best available treatment (1). The pathogenesis of TBM is not well understood, but poor outcomes have been linked to dysregulated intracerebral inflammation (2). Current therapeutic approaches are aimed at killing *M. tb* infecting the brain or controlling the inflammatory response (3, 4). To date, the latter has primarily involved adjunctive corticosteroid therapy and has been associated with increased survival, although how corticosteroids modulate intra-cerebral inflammation to influence outcomes remains uncertain (5).

It is now well established that under ideal conditions the body activates evolutionarily conserved programs to terminate inflammation and promote the repair and regeneration of damaged tissues (6). At the helm of these programs is a recently uncovered genus of mediators produced *via* the stereoselective conversion of essential fatty acids and termed specialized pro-resolving mediators (SPM). These mediators include the arachidonic acid (AA)-derived lipoxins and the n-3 docosapentaenoic acid (DPA)-derived resolvins, protectins and maresins (6). Recent studies demonstrate that SPM via the activation of cognate receptors regulate the phagocytosis and killing of bacteria during infections. They counter regulate the production of pro-inflammatory mediators including cytokines and eicosanoids, and control both leukocyte trafficking and phenotype (7–11). These autacoids also mediate the protective actions of several widely used therapeutics, including atorvastatin, pravastatin and aspirin (7, 12, 13).

We recently tested the hypothesis that the addition of aspirin to standard TBM treatment (anti-tuberculosis drugs and corticosteroids) may further improve outcomes by inhibiting thromboxane A_2_ (TxA_2_) and preventing brain infarcts (a common life-threatening complication of TBM), and by enhancing the resolution of intra-cerebral inflammation through the increased expression of SPMs (14). The trial found that in patients with microbiologically confirmed TBM, aspirin was associated with reduced brain infarcts and/or death in the first 60 days of treatment.

Little is known about the regulation of lipid mediators, and in particular SPM, in the cerebrospinal fluid (CSF) during TBM. Thus, in the present studies we investigated whether CSF lipid mediator concentrations were altered with increasing disease severity. We found that concentrations of a number of SPM families were reduced with increasing disease severity. This was linked with an upregulation of inflammation-initiating eicosanoids, including prostaglandins and cystenyl leukotrienes. Pre-treatment CSF lipid mediator concentrations were also found to be predictive of outcome with distinct lipid mediator profiles obtained in those patients that were alive at the end of the study versus those that died during the study. Finally, we also found that in aspirin-treated patients there was a dose-dependent alteration of the CSF lipid mediator profiles improving the balance between pro-resolving and pro-inflammatory/pro-thrombogenic mediators.

## Results

### Increased disease severity is associated with a reduction in CSF SPM concentrations

In order to determine whether there was a relationship between CSF lipid mediator concentrations and disease severity in TBM we investigated lipid mediator profiles of patients recruited to the Aspirin TBM study (NCT02237365). Here CSF was collected from enrolled patients with known or suspected TBM just prior to the start of treatment in a randomised comparison of aspirin versus placebo as an adjunct to dexamethasone administration for the first 60 days of TBM treatment (14). Disease severity was assessed using the Medical Research Council grade (15) that uses Glasgow coma score and focal neurological deficits to categorise disease severity as mild (grade 1), moderate (grade 2), or severe (grade 3; See Supplemental Table 1 for patient information). Using targeted liquid chromatography-tandem mass spectrometry (LC-MS/MS)-based profiling of CSF we identified mediators from all four major bioactive metabolomes, including the AA-derived lipoxin (LX)s, leukotriene (LT)s and prostaglandin (PG)s and the n-3 DPA, eicosapentaenoic acid (EPA) and docosahexaenoic acid (DHA)-derived resolvin (Rv)s (Supplemental Table 2). In order to gain insights into the relationship between CSF lipid mediator concentrations and disease severity we first grouped the mediators by biological function assessing the differences between pro-resolving mediators (lipoxins, resolvins, protectins and maresins) and pro-inflammatory mediators (leukotrienes, prostaglandins and thromboxane) in patients with moderate or severe disease (MRC grades 2 and 3) when compared with those with mild disease (MRC grade 1)(16)(Figure 1A, Supplemental Table 2). Results of this analysis demonstrated that with increasing disease severity there was a decrease in overall pro-resolving lipid mediator concentrations in the CSF of patients with TBM (Figure 1A). This relationship was coupled with a trend towards an increase in the concentrations of pro-inflammatory mediators in those with more severe disease (Figure 1B). Assessment of the resolution index, that is the ratio between the concentrations of pro-resolving mediators and pro-inflammatory eicosanoids, (16) also demonstrate a reduction in the resolution index, indicative of increased inflammation, in patients with the more severe disease (Figure 1C).

**Figure 1:**
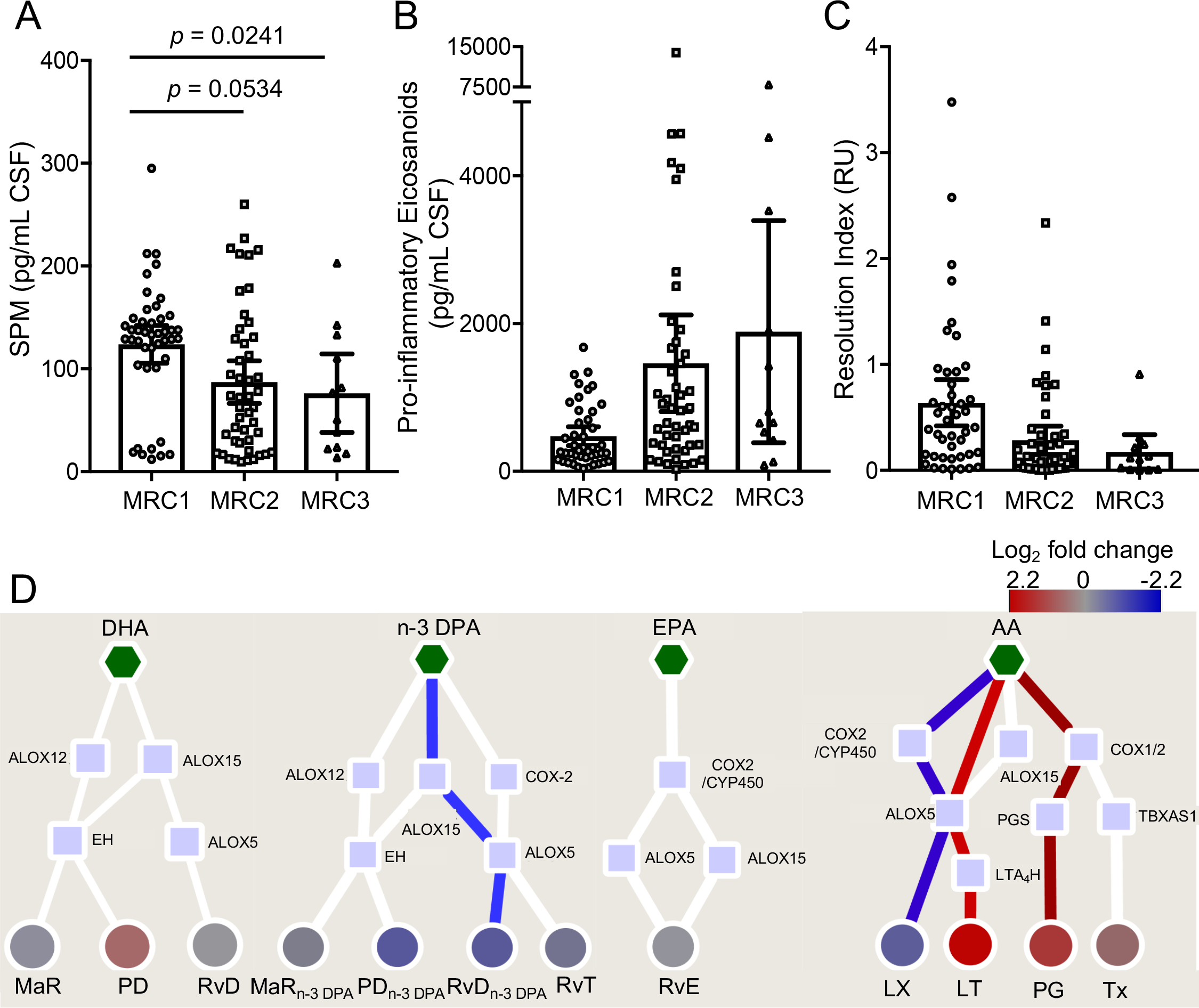
Reduced pre-treatment CSF SPM concentrations are is associated with increased disease severity in TBM. CSF were collected from patients with TBM before the start of treatment and lipid mediators were extracted, identified and quantified using lipid mediator profiling. (A) Sum of pro-resolving mediators (DHA-derived RvD, PD, PCTR, MaR, MCTR; n-3 DPA-derived RvT; RvD_n-3 DPA_, PD_n-3 DPA_, MaR_n-3 DPA_; EPA-derived RvE; AA-derived LX) (B) Sum of pro-inflammatory eicosanoids (AA-derived LT, PG, Tx). (C) Resolution index, the ratio between the sum of pro-resolving mediators and the sum of pro-inflammatory eicosanoids. Statistical differences were determined using Spearmans correlation. Results are mean ± 95% C.I. n = 44 for MRC grade 1; n = 47 for MRC grade 2 and n = 12 for MRC grade 3. (D) Pathway interaction analysis down each of the lipid mediator families. Figure depicts the fold change, expressed as log_2_-fold change, in lipid mediator concentrations between patients with an MRC grade of 1 (MRC1) and those with MRC grade of 2 (MRC2) and MRC grade of 3 (MRC3). Scales represent fold increase or decrease for each mediator family. Mediator families coloured in red or blue represent those families that were found to be significant regulated. Statistical significance between mediator concentrations in patients with MRC1 and those with MRC2 and MRC3 grades was determined using Benjamini-Hochberg multiple testing correction. Red lines depict pathways that are upregulated, in patients with an MRC grade of 2 and 3 when compared with MRC1 patients. Blue lines depict pathways that are downregulated in patients with an MRC grade of 2 and 3 when compared with MRC1 patients. Pentagons depict the distinct essential fatty acids, squares the lipid mediator biosynthetic enzymes and circles the distinct lipid mediator families.DHA = docosahexaenoic acid; n-3 DPA = n-3 docosapentaenoic acid; EPA = eicosapentaenoic acid; AA = arachidonic acid; ALOX = lipoxygenase; COX = cyclooxygenase; CYP450 = Cytochrome P450; LTC_4_S = leukotriene C4 synthase; GSTM = Glutathione S-transferase Mu; LTA_4_H = Leukotriene A_4_ hydrolase; PGS = Prostaglandin synthase; TBXAS1 = thromboxane-A synthase; Rv = resolvins; PD = Protectins; MaR = Maresins; LX = Lipoxins; LT-Leukotrienes; PG = prostaglandins; Tx = Thromboxane.

In order to gain further insights into the mediator pathways that were differentially regulated between these patient groups we next interrogated the biosynthetic pathways for each of the essential fatty acid metabolomes. This demonstrated that in patients with an MRC score of 2 and 3, when compared with patients with an MRC score of 1, there was a significant reduction in two pro-resolving mediator families, the n-3 docosapentaenoic acid-derived resolvins (RvD_n-3 DPA_) and the arachidonic acid-derived lipoxins (LX). This was coupled with a significant increase in the pro-inflammatory arachidonic acid-derived prostaglandins (PG) and leukotrienes (LT; Figure 1D).

We next employed Partial Least Squares Discriminant Analysis (PSL-DA), a regression model that identifies variables that contribute to the separation of experimental groups, to investigate the relationship between lipid mediator profiles (i.e. the concentrations of individual mediators identified in the CSF) and disease severity. This analysis demonstrated that the individual lipid mediator concentrations were markedly different between patients with MRC grades 1, 2 and 3 as demonstrated by the distinct clustering of patients from the different disease severity groups (Figure 2A). Assessment of the variable importance in projection (VIP) scores, which identify the contribution of each mediator in the observed separation between each of the groups, identified 24 mediators that had a VIP score greater than 1, showing that they were differentially regulated depending on disease severity (Figure 2A). Amongst the mediators found to be differentially regulated between these three disease groups were several ALOX5-derived mediators that are involved in coordinating the host response to clear bacterial infections, including RvT4, RvD1 and RvE1 (Figure 2B)(7, 17, 18).

**Figure 2:**
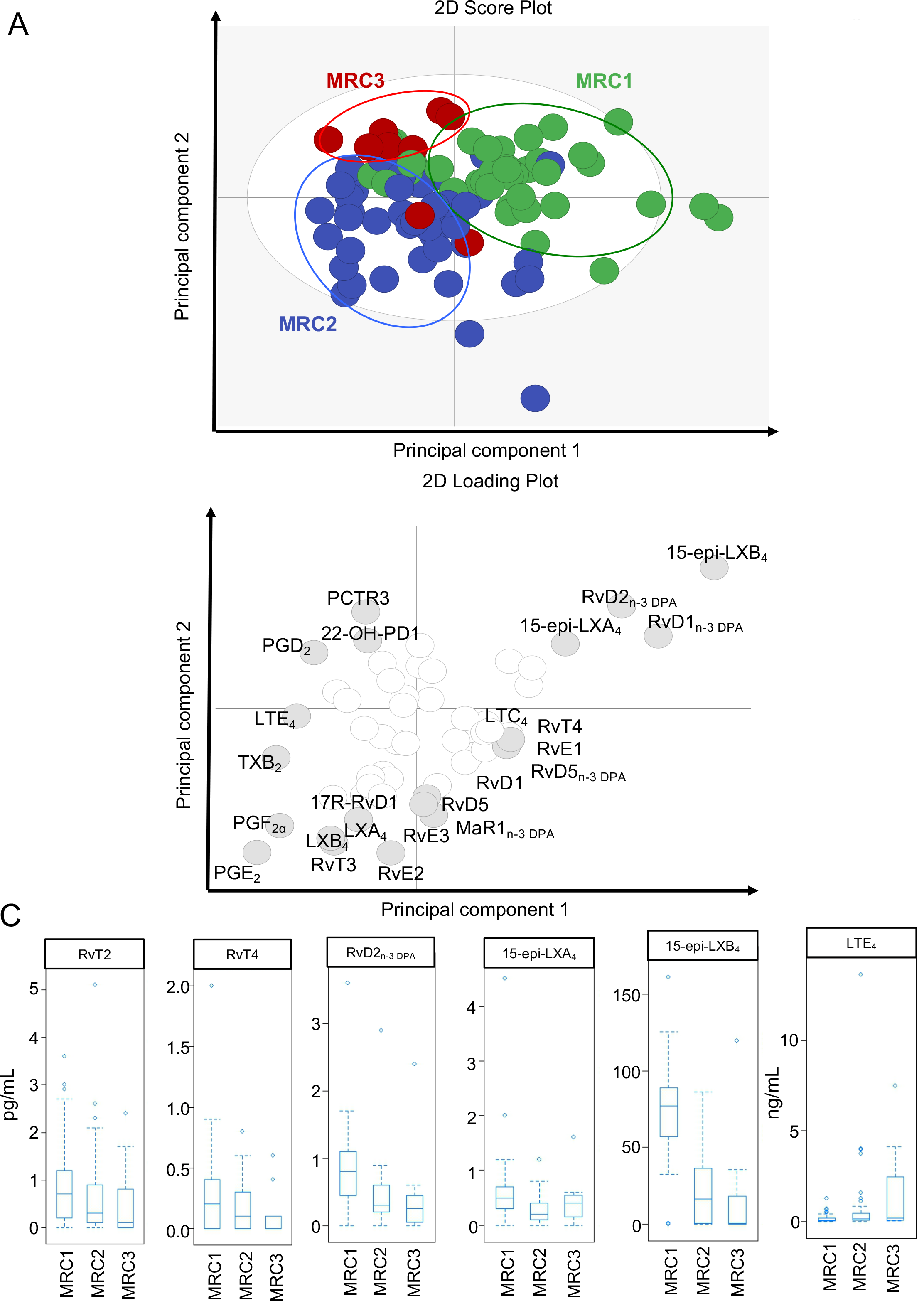
TBM disease severity is linked with a switch in the ALOX5 product profile. CSF were collected from patients with TBM before the start of treatment and lipid mediators were extracted, identified and quantified using lipid mediator profiling. An interaction network was constructed depicting the number of mediators in each of the lipid mediator families that were significantly downregulated (blue) or upregulated (red) in CSF from patients with an MRC grade of 2 and 3 when compared with patients that had an MRC grade of 1. (B) Box-plot of lipid measurement vs. MRC grade of 6 lipid mediators which are significantly associated with disease severity. N = 44 for MRC grade 1; n = 47 for MRC grade 2 and n = 12 for MRC grade 3

Having identified that there was a differential regulation of lipid mediator profiles with increasing disease severity we next assessed the association between distinct lipid mediators and disease severity. There was a significant negative correlation between a select group of pro-resolving mediators, 15-epi-LXB_4_, RvD2_n-3 DPA_, 22-OH-PD1, MaR1 and 15-epi-LXA_4_, and increasing disease severity (Figure 2C and Supplemental Table 2). In addition, we also observed increased concentrations of LTE_4_, the terminal product in the cysteinyl leukotriene biosynthetic metabolome and that was recently found to be also bioactive (19), with increasing disease severity, although the correlation was not statistically significant after correction for multiple testing (Figure 2C, Supplemental Table 2).

To further validate the potential utility of these mediators as biomarkers of disease severity we conducted multiple regression analysis, using Least Absolute Shrinkage and Selection Operator (LASSO) regression analysis. This demonstrated that decreased concentrations of identified pro-resolving mediators, primarily 15-epi-LXB_4_, LXB_4_, PGE_2_, 22-OH-PD1, were associated with increased disease severity (Supplementary Table 3).

Given the role that immune cells play in the biosynthesis of lipid mediators (7, 9), we next investigated whether disease severity influenced CSF leukocyte numbers. Here we found that cell numbers were reduced in those with more severe disease, from 416±53 cells/mm^3^ in patients with an MRC score of 1, to 239±27 cells/mm^3^ and 164±39 cells/mm^3^ in patients with an MRC score of 2 and 3 respectively, although these changes were not statistically significant (p>0.05). We next assessed whether there was an association between CSF leukocyte counts and the concentrations for each of the identified LM families. With adjustment for multiple testing, no significant correlations were found between these parameters (data not shown), suggesting that the observed difference may be due to a differential activation of the leukocyte population.

### Pre-treatment SPM and eicosanoid concentrations in CSF correlate with mortality

Having found SPM concentrations were associated with TBM disease severity we investigated whether CSF lipid mediator concentrations correlated with mortality (See Supplemental Table 4 for patient information). We first assessed the CSF concentrations of SPM and pro-inflammatory eicosanoids, finding that SPM concentrations were significantly reduced in patients that died during the study, while pro-inflammatory mediator concentrations tended to be higher in those who survived, although this did not reach statistical significance (Figure 3A,B and Supplemental Table 5). Of note, comparison of the resolution index in survivors and those who died demonstrated a significantly higher CSF resolution index in survivors (Figure 3C), suggesting that disruption of pro-resolving mediator production and a concomitant increase in pro-inflammatory eicosanoid production are linked with outcome from TBM.

**Figure 3:**
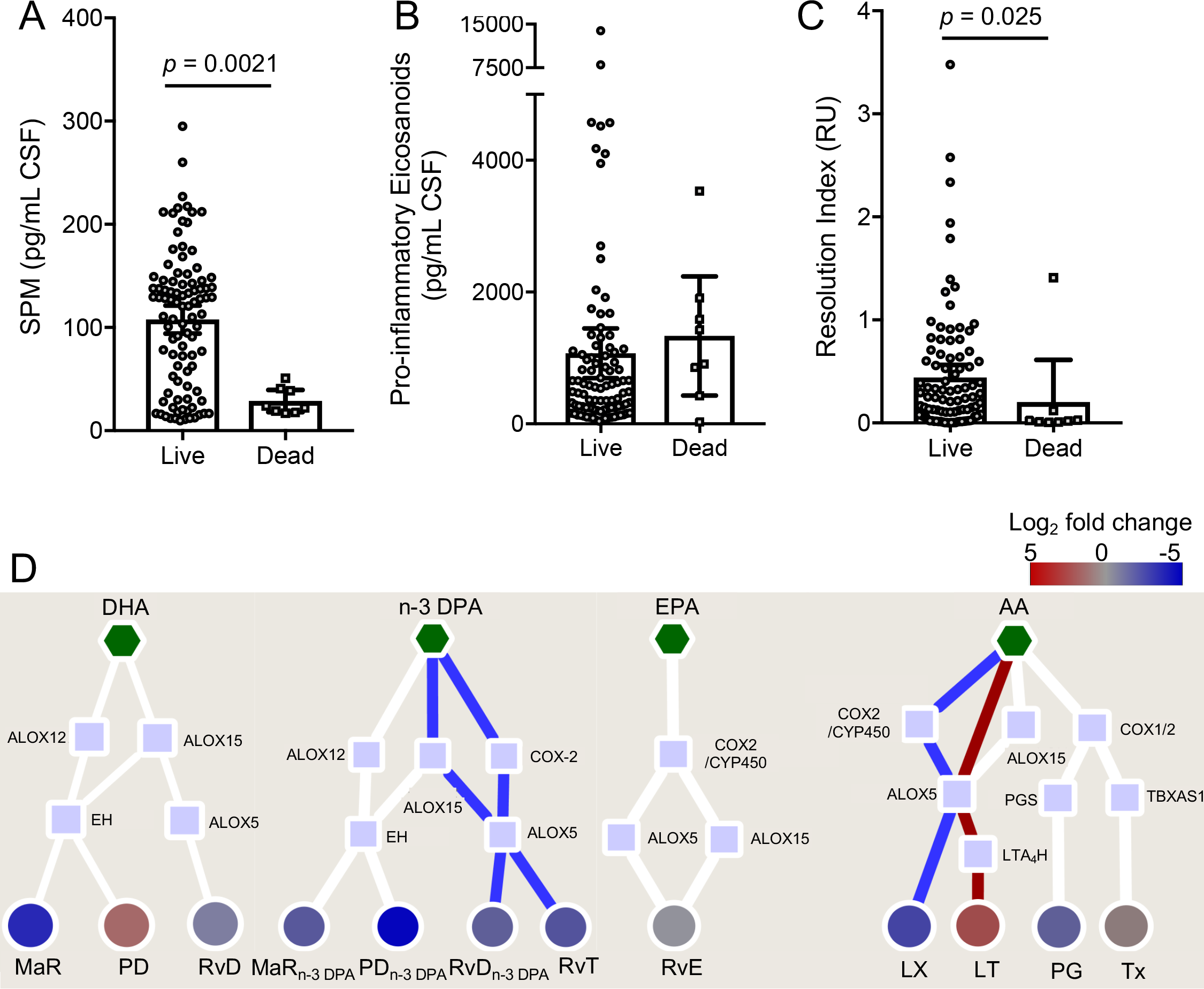
Pre-treatment CSF resolution status is associated with poor outcome from TBM. CSF was collected before the start of treatment and lipid mediators identified and quantified using lipid mediator profiling. Sum of pro-resolving mediators; (B) Sum of pro-inflammatory eicosanoids; (C) Resolution index. Statistical analysis was conducted using one-way ANOVA with Dunnet post-hoc test. Results are mean ± 95% C.I. n = 95 survivors and 8 non-survivors. (D) Interaction networks were constructed to comparing the pre-treatment CSF lipid mediator profiles from patients that died during the 80-day trial period to those that survived. Scales represent fold increase or decrease for each mediator family. Mediator families coloured in red or blue represent those families that were found to be significant regulated. Statistical significance was determined using Benjamini-Hochberg multiple testing correction. Results are representative of n = 8 patients. Red lines depict pathways that are upregulated in non-survivors. Blue lines depict pathways that are downregulated in non-survivors. Pentagons depict the distinct essential fatty acids, squares the lipid mediator biosynthetic enzymes and circles the distinct lipid mediator families.

We next conducted lipid mediator biosynthetic pathway analysis to evaluate which pathways are contributing to the observed differences in lipid mediator concentrations. This demonstrated that there was a downregulation in the expression of 13-series resolvins (RvT), RvD_n-3 DPA_, and LX with an upregulation in the pro-inflammatory LT in non-survivors when compared with survivors (Figure 3D).

Orthogonal PLS-DA (OPLS-DA) analysis of CSF lipid mediator profiles provided further support for the differences in lipid mediator concentrations between those who died and those who survived, with CSF lipid mediator concentrations from each of these patient groups giving two distinct clusters (Figure 4A). This separation between the two groups was linked with a differential regulation of 18 lipid mediators, which gave a VIP score greater than 1 and included MaR2, RvT2 and 15-epi-LXB_4_. Of note, RvT2 and 15-epi-LXB_4_ activate the innate immune response to clear bacterial infections and counter-regulate the production of pro-inflammatory mediators (6, 7). Statistical assessment of mediators found to be differentially regulated demonstrated that the concentrations of MaR2, RvT2 and 15-epi-LXB_4_ were significantly lower in non-survivors when compared with survivors (Supplemental Table 5 and Figure 4B). Out of these mediators 15-epi-LXB_4_ was found to be significantly reduced after multiple correction. This lipid mediator was also a strong predictor of mortality in TBM as demonstrated by LASSO regression analysis (Supplemental Table 6).

**Figure 4:**
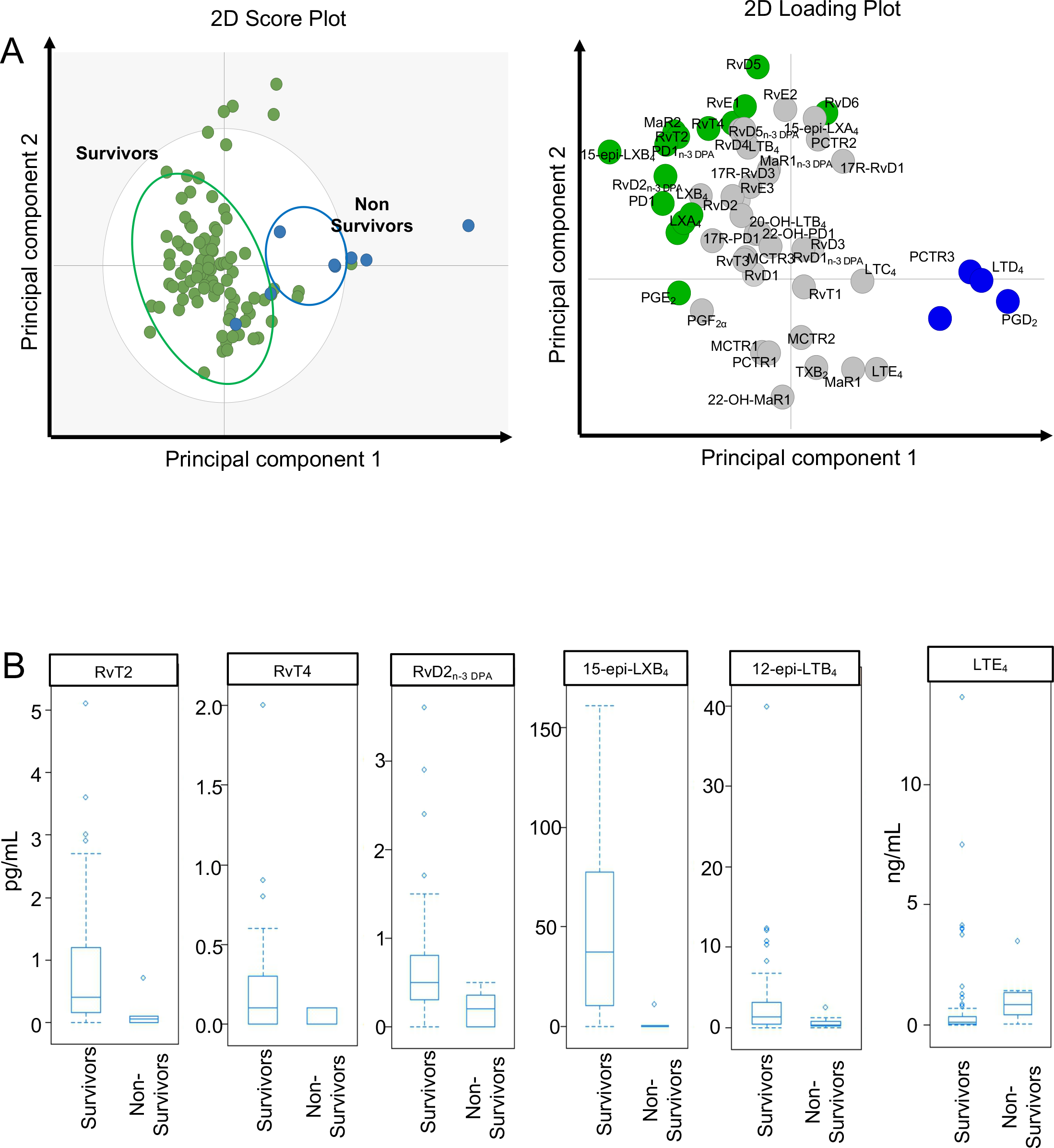
Death from TBM is associated with a downregulation of CSF ALOX5-derived SPM and an upregulation of ALOX5 derived LTE_4_. (A) CSF lipid mediator profiles obtained from patients that died during the 80-day dration of the study (non-survivors) were compared with those that were alive at the end of the study (survivors) using OPLS-DA (left panel) Score plot. (right panel) loading plot. (B) Box-plot of lipid measurement vs. 80-day mortality outcome of 6 lipid mediators which are significant associated with mortality. n = 95 survivors and 8 non-survivors

### Aspirin administration upregulates CSF concentrations of select pro-resolving mediators during TBM

We recently reported that treatment with aspirin, dexamethasone and anti-tuberculosis drugs was associated with reduction in new brain infarcts and deaths within 60 days in patients with microbiologically confirmed TBM (14). Therefore, we investigated the impact of aspirin on CSF lipid mediator pathways in this patient sub-population. We compared lipid mediator profiles obtained after 30 days of aspirin (1000mg/day or 81mg/day) or placebo (Supplementary Table 7). Assessment of overall lipid mediators by function demonstrated a decrease in the concentrations of proinflammatory eicosanoids and an overall increase in the resolution index of patients given 1000mg of aspirin when compared with those receiving placebo (Figure 5A-C). However, these changes did not reach statistical significance. Pathway analysis of lipid mediator families identified in the CSF of these patients demonstrated a trend towards the upregulation of a number of pro-resolving families that included the RvD_n-3 DPA_ and MaR_n-3 DPA_ in patients receiving 1000mg and 81mg aspirin respectively, although these changes did not reach statistical significance (n= 28 for patients in 81mg aspirin group, n = 27 patients in 1000mg aspirin group and n= 34 patients in placebo group).

To further evaluate the regulation of CSF lipid mediator concentrations by aspirin we conducted OPLS-DA. This analysis demonstrated that while day 30 CSF lipid mediator profiles between the 81mg and placebo groups were not markedly different, lipid mediator profiles from patients given 1000mg aspirin gave distinct clusters to that from patients given placebo (Figure 5D,E). This separation between the two patient groups was linked with a differential regulation of 18 mediators from all the four bioactive metabolomes, including RvT4 and TxB_2_, the inactive breakdown product of the prothrombotic TxA_2_ (20)(Figure 5E). Statistical assessment of their concentrations in CSF demonstrated that after 30 days TxB_2_ was reduced with both 81 mg and 1000mg of aspirin, reaching statistical significance in those given 1000mg after correcting for multiple testing (Supplemental Table 8,9).

**Figure 5.**
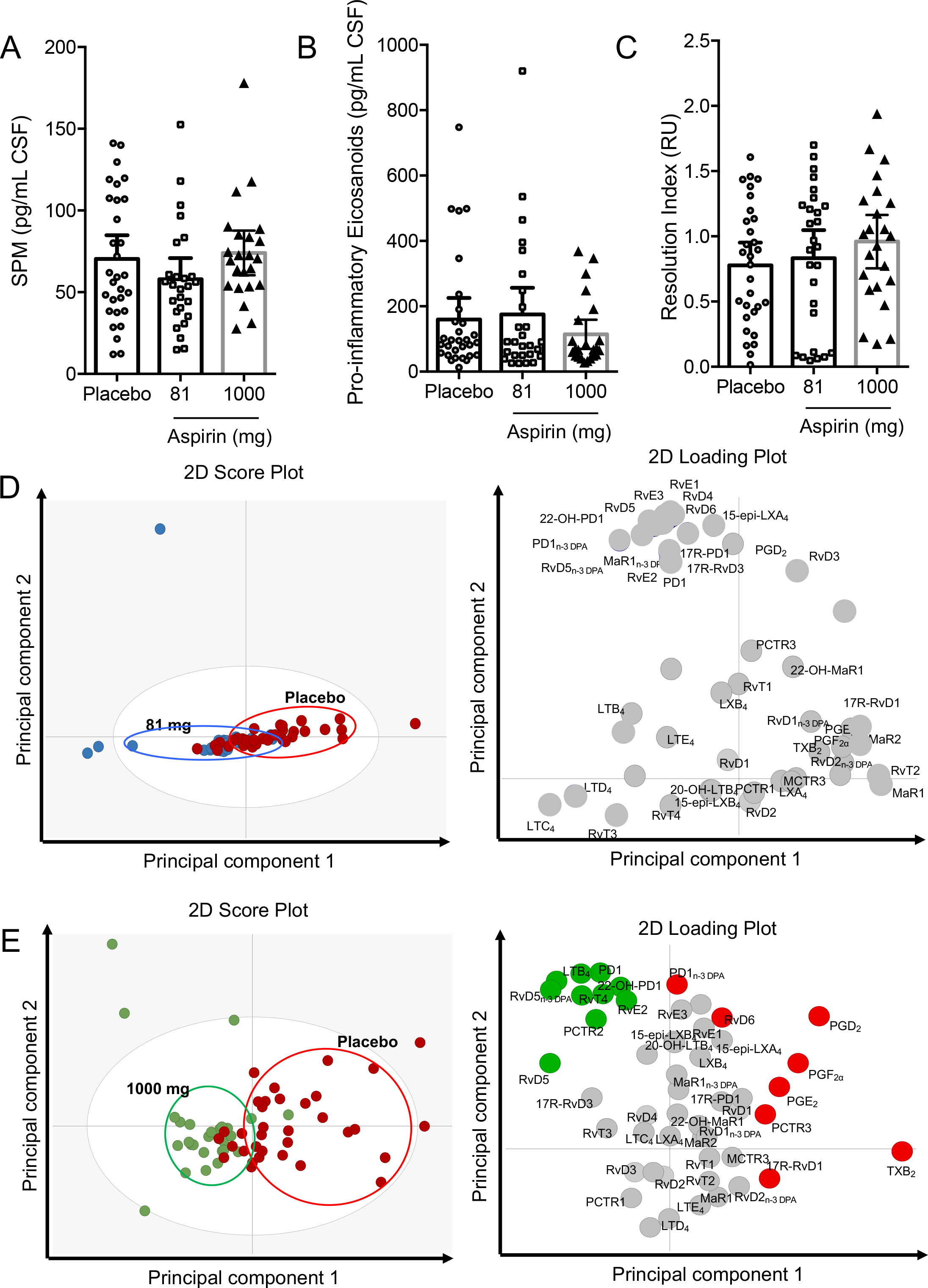
Aspirin 1000mg/day increases CSF resolution status and reduces CSF TxB_2_ concentrations after 30 days of treatment. CSF fluids were collected 30 days after administration of 81mg, 1000mg aspirin per day or placebo. Lipid mediators were extracted, identified, quantified using lipid mediator profiling. (A) cumulative pro-resolving mediator concentrations (B) cumulative pro-inflammatory eicosanoid concentrations (C) resolution index. (D,E) OPLS-DA of lipid mediator profiles from patients given (D) 81mg (E) 1000mg aspirin per day in comparison to those receiving placebo. (*Left panel*) score plot; (*right panel*) loading plot. Mediators with a VIP score > 1 are identified in red (placebo) green (1000mg aspirin) or blue (81mg aspirin) circles that denote the association with the placebo and aspirin group. Results for A-C are mean ± 95% C.I. n= 28 for patients in 81mg aspirin group and n = 27 patients in 1000mg aspirin group and n= 34 patients in placebo group.

## Discussion

In the present study we investigated the regulation of CSF lipid mediator profiles before and during the treatment of adults with TBM. We found that pre-treatment disease severity was associated with the concentrations of both inflammatory and pro-resolving mediators, with more severe disease linked to lower SPM concentrations and increased concentrations of immunosuppressive, vasoconstrictive and nociceptive eicosanoids. Pre-treatment SPM concentrations were also associated with 80-day mortality, with survivors having higher concentrations of several SPM families, including the ALOX5-derived Rvs and LXs, compared to those who died. Aspirin co-administration with dexamethasone was observed to increase the CSF resolution index after 30 days of treatment, and decrease TxB_2_ concentrations.

Recent decades have seen significant advances in our ability to manage patients with TBM, however, morbidity and mortality remain high (1). This is at least part due to our limited understanding of the underlying mechanisms that perpetuate inflammation within the CNS leading to disability and ultimately death despite the best available treatment. In the present studies we investigated the relationship between disease severity and local mediator biosynthesis. Results from these analysis demonstrate that even prior to the initiation of treatment patients that died during the course of the study presented with a profound dysregulation in both protective and inflammatory mediator pathways.

Lipid mediator biosynthesis is a tightly coordinated process where essential fatty acids are sequentially oxygenated, primarily by lipoxygenase and cyclooxygenase enzymes, to produce stereochemically defined and structurally unique products (6, 21). Regulation of these biosynthetic enzymes occurs at both a transcriptional/translational level as well as via post-translational modifications. Recent studies demonstrate that nitric oxide synthase promotes the S-nitrosylation of COX-2 increasing its catalytic activity and upregulating the production of protective mediators including PGI_2_ (22) and RvT (7). On the other hand, the calcium-calmodulin-dependent protein kinase II - p38 mediated phosphorylation of ALOX5 leads to a switch in the product profile of the enzyme from SPM production to the formation of leukotrienes (23). The observation that with increasing disease severity there is a downregulation of ALOX5-derived SPM and a concomitant upregulation in cysteinyl leukotriene production suggest that as the disease progresses there is an increase in ALOX5 phosphorylation that leads to an alteration in the product profile of the enzyme.

Leukocytes play an important role in the biosynthesis of lipid mediators, with their product profile reflecting their activation status (23–25). Different macrophage subsets, for example, display distinct lipid mediator profiles with monocyte-derived-macrophages skewed towards a classic phenotype expressing higher amounts of pro-inflammatory eicosanoids, whereas cells with an alternatively activated phenotype display higher concentrations of pro-resolving mediators (23–25). This shift is also linked with a differential expression of both lipid mediator biosynthetic enzymes as well as the phosphorylation status of ALOX5 (23, 25, 26). In the present study we found a decrease in the number of leukocytes in the CSF. However there was no correlation between leukocyte numbers and lipid mediator concentrations suggesting that regulation of enzyme activity, possibly reflecting a shift in leukocyte phenotype or population, is responsible for the altered lipid mediator concentrations.

Studies conducted by Vane and colleagues demonstrate that aspirin inhibits the production of prostaglandins and thromboxane, a mechanism that is dependent on the acetylation of COX-1 and COX-2 (27). Later investigations by Serhan and colleagues found that acetylation of COX-2 also led to a switch in the catalytic activity of the enzyme, from the production of PGG_2_ to the formation of epimeric forms of resolvins, lipoxins and protectins (28, 29). In the preset study we did not observe significant increases in these epimeric forms of the SPM in the CSF of TBM patients treated with aspirin. This may be because the present study was not adequately powered for this analysis or because at the interval tested, i.e. 30 days post initiation of treatment, the biosynthetic pathways leading to the formation of these molecules were downregulated. Future studies powered to interrogate this question will need to establish whether daily regulation of high dose aspirin downregulates these pathways in the CSF. Of note, we found high dose aspirin administration reduced TxB2 concentrations and was linked with improved outcomes.

In summary, the present findings uncover novel mechanisms in the pathophysiology of TBM that may have relevance to all forms of tuberculosis. Disease severity is associated with an alteration in lipid mediator expression and a dysregulation in the production of several SPM pathways; a mechanism that is also linked with a poor prognosis. This dysregulation was at least in part restored by aspirin, lending support to the hypothesis that TBM arises from a failure of the host to engage resolution mechanisms and offering novel treatment strategies.

## Materials and Methods

### Clinical study

The patients in the current study were enrolled in a clinical trial of adjunctive aspirin, the design and conduct of which have been previously described (14). Briefly, we conducted a parallel group, double blind, randomised, placebo-controlled trial in HIV-uninfected adults with TBM to assess the safety and efficacy of either 81mg or 1000mg aspirin daily for the first 60 days of treatment with standard anti-tuberculosis drugs and dexamethasone. The trial enrolled in-patients at the Hospital for Tropical Diseases, a 550-bed tertiary referral hospital in Ho Chi Minh City, Vietnam, and was approved by the Oxford Tropical Research Ethics Committee and the Institutional Review Board of the Hospital for Tropical Diseases and the Ethical Committee of the Ministry of Health, Vietnam. Adults (≥ 18 years old) with suspected TBM (at least 5 days of meningitis symptoms, nuchal rigidity, and CSF abnormalities) and a negative HIV test were eligible to enter the trial. Written informed consent to participate in the study was obtained from all participants or from their relatives if the participant could not provide consent due to incapacity.

Lumbar puncture was performed before the start of treatment and on days 30 and 60 as per normal clinical care with CSF archived at −80°C until later testing. Clinical progress and neurological and drug-related adverse events were assessed daily until discharge from hospital and monthly thereafter until 8 months, when a final clinical assessment was made.

#### Targeted lipid mediator profiling

All samples were extracted using solid-phase extraction columns as in (30). Prior to sample extraction, deuterated internal standards, representing each region in the chromatographic analysis (500 pg each) were added to facilitate quantification in 4V of cold methanol. Samples were kept at −20°C for a minimum of 45 min to allow protein precipitation. Supernatants were subjected to solid phase extraction, methyl formate and methanol fractions were collected, brought to dryness and suspended in phase (methanol/water, 1:1, vol/vol) for injection on a Shimadzu LC-20AD HPLC and a Shimadzu SIL-20AC autoinjector, paired with a QTrap 6500 plus (Sciex). For identification and quantitation of products eluted in the methyl formate, an Agilent Poroshell 120 EC-C18 column (100 mm × 4.6 mm × 2.7 μm) was kept at 50°C and mediators were eluted using a mobile phase consisting of methanol-water-acetic acid of 20:80:0.01 (vol/vol/vol) that was ramped to 50:50:0.01 (vol/vol/vol) over 0.5 min and then to 80:20:0.01 (vol/vol/vol) from 2 min to 11 min, maintained till 14.5 min and then rapidly ramped to 98:2:0.01 (vol/vol/vol) for the next 0.1 min. This was subsequently maintained at 98:2:0.01 (vol/vol/vol) for 5.4 min, and the flow rate was maintained at 0.5 ml/min. QTrap 6500+ was operated in negative ionization mode using a multiple reaction monitoring method as in (30).

In the analysis of peptide-lipid conjugated mediators eluted in the methanol fraction, an Agilent Poroshell 120 EC-C18 column (100 mm × 4.6 mm × 2.7 μm) was kept at 50°C and mediators were eluted using a mobile phase consisting of methanol-water-acetic acid at 55:45:0.1 (vol:vol:vol) over 5 min, that was ramped to 80:20:0.1 (vol:vol:vol) for 2 min, maintained at 80:20:0.1 (vol:vol:vol) for the next 3 min and ramped to 98:2:0.1 (vol:vol:vol) over 3 min. This was kept at 98:2:0.1 (vol:vol:vol) for 3 min. A flow rate of 0.65 ml/min was used throughout the experiment. QTrap 6500+ was operated in positive ionization mode using scheduled multiple reaction monitoring (MRM) coupled with information-dependent acquisition and enhanced product ion scan (30).

Each LM was identified using established criteria including matching retention time to synthetic and authentic materials and at least 6 diagnostic ions (30). Calibration curves were obtained for each using synthetic compound mixtures at 0.78, 1.56, 3.12, 6.25, 12.5, 25, 50, 100, and 200 pg that gave linear calibration curves with an r^2^ values of 0.98–0.99.

### Statistical analysis

We performed all statistical analyses and data derivation and prepared the manuscript in R (31), Prism 8 and Microsoft Excel. Results are expressed as mean and 95% confidence interval or interquartile range as indicated in the figures and tables. Summary tables of the baseline characteristics of the study population with respect to disease severity, mortality outcome and treatment allocation as median (IQR) for continuous data and n (%) for categorical data were provided in the supplemental document.

To assess the overall balance between pro-inflammatory mediators and pro-resolving mediators in the CSF we combined the concentrations of pro-resolving mediators, combining the concentrations of the DHA-derived RvD (RvD1, RvD2, RvD3, RvD4, RvD5, RvD6, 17R-RvD1 and 17R-RvD3) PD (PD1, 10S,17S-diHDHA, 17R-PD1 and 22-OH-PD1) PCTRs (PCTR1, PCTR2 and PCTR3) and MaR (MaR1,7S, 14S-diHDHA, MaR2, 4S, 14S-diHDHA and 22-OH-MaR1) MCTRs (MCTR1, MCTR2 and MCTR3), the n-3 DPA derived RvT (RvT1, RvT2, RvT3 and RvT4), RvD_n-3 DPA_ (RvD1_n-3 DPA_, RvD2_n-3 DPA_ and RvD5_n-3 DPA_), PD_n-3 DPA_ (PD1_n-3 DPA_ and 10S, 17S-diHDPA) and MaR_n-3 DPA_ (MaR1_n-3 DPA_ and 7S, 14S-diHDPA), the EPA-derived RvE (RvE1, RvE2 and RvE3) and the AA-derived LX (LXA_4_, LXB_4_, 5S, 15S-diHETE, 15R-LXA_4_ and 15R-LXB_4_). Separately we combined the concentrations of pro-inflammatory eicosanoids: AA-derived LT (LTB_4_, 5S, 12S-diHETE, 12-epi-LTB_4_, 6-trans, 12-epi-LTB_4_ and 20-OH-LTB_4_), cysLT (LTC_4_, LTD_4_ and LTE_4_), PG (PGD_2_, PGE_2_ and PGF_2α_) and Tx (TxB_2_). The resolution index was obtained by dividing the overall concentration of pro-resolving mediators by the overall concentrations of pro-inflammatory eicosanoids. Investigations into the *flux down each of the mediator families* was conducted by determining the fold change in the concentration of the different mediator families indicated above, between the control groups (ie MRC1 or survivors) and test groups (ie MRC2 + MRC3 or non-survivors respectively).

Statistical differences between the concentrations (expressed as the log2-fold change) of the mediators in each analysis group was determined using Hochberg Multiple Testing Correction or Benjamini-Hochberg Multiple Testing Correction as indicated. Lipid mediator networks were constructed using Cytoscape 3.7.1. and the pathways that were statistically up or down regulated were denoted using red and blue lines respectively.

Investigators were not blinded to group allocation or outcome assessment. The criterion for statistical significance was *p* ≤ 0.05. OPLS-DA and PLS-DA (32) were performed using SIMCA 14.1 software (Umetrics, Umea, Sweden) following mean centering and unit variance scaling of LM levels. PLS-DA is based on a linear multivariate model that identifies variables that contribute to class separation of observations (MRC scores, survivors/non-survivors; Placebo/aspirin groups) on the basis of their variables (LM levels). During classification, observations were projected onto their respective class model. The score plot illustrates the systematic clusters among the observations (closer plots presenting higher similarity in the data matrix). Loading plot interpretation identified the variables with the best discriminatory power (Variable Importance in Projection greater than 1) that were associated with the distinct intervals and contributed to the tight clusters observed in the Score plot. Comparisons of lipid mediators by disease severity (MRC grade) or mortality outcome were based on the Spearman-test for the test for trend of ordinary groups for non-normal continuous outcomes. The test were implemented in R package “Comparegroup”. In addition, to identify which lipid mediators are the strong predictor for disease severity and mortality outcomes in TBM, we performed a Lasso regression model in which the disease severity or mortality as the outcome, and all the lipid mediators as the covariates. The lipid mediators which were likely chosen by the model demonstrated as the strong predictors of the disease severity or mortality in TBM.

The reduction of lipid mediator was computed as the difference between the two timepoints from baseline and day 30. The reduction was compared between treatment arms based on a simple linear regression. In this model, the lipid mediator reduction is the outcome and treatment arm is the main covariate. To increase the power for the analysis, the model was also adjusted for the baseline lipid mediator. For all the comparison in this study, we performed multiplicity adjustments of p-values for all comparisons based on Hochberg Multiple Testing Correction, as implemented in R package “stats”.

## Acknowledgment

We thank all the patients who took part in the study and the ward nurses and doctors who cared for them during the trial. We also thank Prof Lalita Ramakrishnan (University of Cambridge, UK) and Prof Paul Edelstein, (University of Pennsylvania, USA) for insightful comments and critical review of the manuscript. JD received funding from the European Research Council (ERC) under the European Union’s Horizon 2020 research and innovation programme (grant no: 677542) and the Barts Charity (grant no: MGU0343). JD is also supported by a Sir Henry Dale Fellowship jointly funded by the Wellcome Trust and the Royal Society (grant 107613/Z/15/Z). GT is supported by a Wellcome Trust Investigator award (grant 110179/Z/15/Z) and GT, NTTH, THP and NTHM are supported by Wellcome Trust core award to the Vietnam Asia programme (grant 106680/Z/14/Z).

## Author Contributions

THP, NTHM recruited and cared for the patients; LTHN and HHT helped to analyse the data; NTTT helped design the study and coordinated the laboratory assessments in Vietnam; GT, RAC, LL, EAGC and JD carried out experiments and analysed data; All authors contributed to manuscript preparation; GT and JD conceived overall research plan.

## References

1. Wilkinson RJ, et al. (2017) Tuberculous meningitis. Nat Rev Neurol 13(10):581–598.

2. Thuong NTT, et al. (2017) Leukotriene A4 Hydrolase Genotype and HIV Infection Influence Intracerebral Inflammation and Survival From Tuberculous Meningitis. J Infect Dis 215(7):1020–1028.

3. Ruslami R, et al. (2013) Intensified regimen containing rifampicin and moxifloxacin for tuberculous meningitis: an open-label, randomised controlled phase 2 trial. Lancet Infect Dis 13(1):27–35.

4. Heemskerk AD, et al. (2016) Intensified Antituberculosis Therapy in Adults with Tuberculous Meningitis. N Engl J Med 374(2):124–134.

5. Prasad K, Singh MB, & Ryan H (2016) Corticosteroids for managing tuberculous meningitis. Cochrane Database Syst Rev 4:CD002244.

6. Dalli J & Serhan CN (2017) Immunoresolvents signaling molecules at intersection between the brain and immune system. Curr Opin Immunol 50: 48–54.

7. Dalli J, Chiang N, & Serhan CN (2015) Elucidation of novel 13-series resolvins that increase with atorvastatin and clear infections. Nat Med 21(9):1071–1075.

8. Chiang N, Dalli J, Colas RA, & Serhan CN (2015) Identification of resolvin D2 receptor mediating resolution of infections and organ protection. J Exp Med 212(8):1203–1217.

9. Gao Y, et al. (2015) Female-Specific Downregulation of Tissue Polymorphonuclear Neutrophils Drives Impaired Regulatory T Cell and Amplified Effector T Cell Responses in Autoimmune Dry Eye Disease. J Immunol 195(7):3086–3099.

10. Fredman G, et al. (2016) An imbalance between specialized pro-resolving lipid mediators and pro-inflammatory leukotrienes promotes instability of atherosclerotic plaques. Nat Commun 7:12859.

11. El Kebir D, Gjorstrup P, & Filep JG (2012) Resolvin E1 promotes phagocytosis-induced neutrophil apoptosis and accelerates resolution of pulmonary inflammation. Proc Natl Acad Sci U S A 109(37):14983–14988.

12. Maderna P & Godson C (2009) Lipoxins: resolutionary road. Br J Pharmacol 158(4):947–959.

13. Chiang N, Hurwitz S, Ridker PM, & Serhan CN (2006) Aspirin has a gender-dependent impact on antiinflammatory 15-epi-lipoxin A4 formation: a randomized human trial. Arterioscler Thromb Vasc Biol 26(2):e14–17.

14. Mai NT, et al. (2018) A randomised double blind placebo controlled phase 2 trial of adjunctive aspirin for tuberculous meningitis in HIV-uninfected adults. Elife 7.

15. Anonymous (1948) STREPTOMYCIN treatment of tuberculous meningitis. Lancet 1(6503):582–596.

16. Colas RA, et al. (2018) Impaired Production and Diurnal Regulation of Vascular RvDn-3 DPA Increases Systemic Inflammation and Cardiovascular Disease. Circ Res.

17. Chiang N, et al. (2012) Infection regulates pro-resolving mediators that lower antibiotic requirements. Nature 484(7395):524–528.

18. Oh SF, Pillai PS, Recchiuti A, Yang R, & Serhan CN (2011) Pro-resolving actions and stereoselective biosynthesis of 18S E-series resolvins in human leukocytes and murine inflammation. J Clin Invest 121(2):569–581.

19. Bankova LG, et al. (2016) Leukotriene E4 elicits respiratory epithelial cell mucin release through the G-protein-coupled receptor, GPR99. Proc Natl Acad Sci U S A 113(22):6242–6247.

20. Hamberg M, Svensson J, & Samuelsson B (1975) Thromboxanes: a new group of biologically active compounds derived from prostaglandin endoperoxides. Proc Natl Acad Sci U S A 72(8):2994–2998.

21. Samuelsson B (2012) Role of basic science in the development of new medicines: examples from the eicosanoid field. J Biol Chem 287(13):10070–10080.

22. Atar S, et al. (2006) Atorvastatin-induced cardioprotection is mediated by increasing inducible nitric oxide synthase and consequent S-nitrosylation of cyclooxygenase-2. Am J Physiol Heart Circ Physiol 290(5):H1960–1968.

23. Fredman G, et al. (2014) Resolvin D1 limits 5-lipoxygenase nuclear localization and leukotriene B4 synthesis by inhibiting a calcium-activated kinase pathway. Proc Natl Acad Sci U S A 111(40):14530–14535.

24. Dalli J & Serhan CN (2012) Specific lipid mediator signatures of human phagocytes: microparticles stimulate macrophage efferocytosis and pro-resolving mediators. Blood 120(15):e60–72.

25. Werz O, et al. (2018) Human macrophages differentially produce specific resolvin or leukotriene signals that depend on bacterial pathogenicity. Nat Commun 9(1):59.

26. Pistorius K, et al. (2018) PDn-3 DPA Pathway Regulates Human Monocyte Differentiation and Macrophage Function. Cell Chem Biol 25(6):749–760 e749.

27. Vane JR, Flower RJ, & Botting RM (1990) History of aspirin and its mechanism of action. Stroke 21(12 Suppl):IV12–23.

28. Claria J & Serhan CN (1995) Aspirin triggers previously undescribed bioactive eicosanoids by human endothelial cell-leukocyte interactions. Proc Natl Acad Sci U S A 92(21):9475–9479.

29. Serhan CN, et al. (2002) Resolvins: a family of bioactive products of omega-3 fatty acid transformation circuits initiated by aspirin treatment that counter proinflammation signals. J Exp Med 196(8):1025–1037.

30. Dalli J, Colas RA, Walker ME, & Serhan CN (2018) Lipid Mediator Metabolomics Via LC-MS/MS Profiling and Analysis. Methods Mol Biol 1730: 59–72.

31. R Development Core Team (2016) R: A language and environment for statistical computing.).

32. Janes KA & Yaffe MB (2006) Data-driven modelling of signal-transduction networks. Nat Rev Mol Cell Biol 7(11):820–828.

