## Supplemental tables for "Pro-Resolving Mediator Profiles And 5-Lipoxygenase Activity In Cerebrospinal Fluid Correlate with Disease Severity and Outcome in Adults with Tuberculous Meningitis"

* Equal first authorship

[$] Equal senior authorship

[+]Corresponding author: Dr Jesmond Dalli Ph.D, William Harvey Research Institute, John Vane Science Centre, Charterhouse Square, London. EC1M 6BQ., Tel: +44 (0) 207 882 8263

**Short Title:** Relationships between SPM levels and outcomes in Tuberculous meningitis

**Keywords:** Tuberculous meningitis, Resolution, essential fatty acids, eicosanoids, aspirin

### Supplemental Tables

#### Supplemental Table 1: Summary data set MRC1, MRC2, MRC3

| Characteristic | n | Summary statistic #MRC1 (N=44) | n | Summary statistic MRC2 (N=47) | n | Summary statistic MRC3 (N=12) |
|---|---|---|---|---|---|---|
| Gender – no. (%) | 44 | | 46 | | 12 | |
| - Male | | 22/44 (50%) | | 34/46 (74%) | | 8/12 (67%) |
| - Female | | 22/44 (50%) | | 12/46 (26%) | | 4/12 (33%) |
| Age (years) – median (IQR) | 44 | 38.00(27.75,50.25) | 46 | 42.50(32.25,49.00) | 12 | 40.00(37.25,45.50) |
| Weight (kg) – median (IQR) | 44 | 52.00(45.00,61.25) | 46 | 50.00(46.00,58.00) | 12 | 55.75(45.38,60.50) |
| Glasgow coma score – median (IQR) | 44 | 15.00(15.00,15.00) | 47 | 13.00(12.00,14.00) | 12 | 8.50(8.00,10.00) |
| *Diagnostic category – no. (%) | 44 | | 47 | | 12 | |
| - definite TBM | | 33/44 (75%) | | 40/47 (85%) | | 8/12 (67%) |
| - possible TBM | | 9/44 (20%) | | 5/47 (11%) | | 1/12 (8%) |
| - probable TBM | | 2/44 (5%) | | 2/47 (4%) | | 3/12 (25%) |
| Treatment arm – no. (%) | 44 | | 47 | | 12 | |
| - Aspirin_1000mg | | 15/44 (34%) | | 13/47 (28%) | | 4/12 (33%) |
| - Aspirin_81mg | | 14/44 (32%) | | 16/47 (34%) | | 5/12 (42%) |
| - Placebo | | 15/44 (34%) | | 18/47 (38%) | | 3/12 (25%) |

All summary statistics are absolute counts (%) for categorical variables and median (inter-quartile range = IQR) for continuous data. n refers to the number of patients with non-missing data for the corresponding variable.
#MRC denotes modified British Medical Research Council criteria. MRC1 indicates a GCS of 15 with no neurologic signs (baseline), MRC2 a score of 11 to 14 (or 15 with focal neurologic signs), and MRC3 a score of 10 or less.
*Diagnostic categories were assigned according to the consensus case definition (Marais et al.; The Lancet Infectious diseases. 2010; **10**(11): 803-12).

### Supplemental Table 2: Comparison of lipid mediators by disease severity (MRC grade)

| Lipid Mediators | | 1 (N=44) | 2 (N=47) | 3 (N=12) | P trend | Adjusted P |
|---|---|---|---|---|---|---|
| **DHA metabolome** | | | | | | |
| RvD1 | Negative* | 27 (61.4%) | 30 (63.8%) | 7 (58.3%) | 0.972 | 0.993 |
| | Positive | 17 (38.6%) | 17 (36.2%) | 5 (41.7%) | | |
| RvD2 | Negative | 27 (61.4%) | 33 (70.2%) | 7 (58.3%) | 0.802 | 0.925 |
| | Positive | 17 (38.6%) | 14 (29.8%) | 5 (41.7%) | | |
| RvD3 | Negative | 33 (75.0%) | 31 (66.0%) | 10 (83.3%) | 0.997 | 0.998 |
| | Positive | 11 (25.0%) | 16 (34.0%) | 2 (16.7%) | | |
| RvD4 | | 0.30 [0.10;0.43] | 0.20 [0.10;0.55] | 0.20 [0.00;0.40] | 0.252 | 0.492 |
| RvD5 | | 0.70 [0.20;1.07] | 0.40 [0.15;1.00] | 0.25 [0.00;0.80] | 0.121 | 0.356 |
| RvD6 | Negative | 31 (70.5%) | 30 (63.8%) | 7 (58.3%) | 0.374 | 0.577 |
| | Positive | 13 (29.5%) | 17 (36.2%) | 5 (41.7%) | | |
| 17R-RvD1 | Negative | 26 (59.1%) | 23 (48.9%) | 3 (25.0%) | 0.045** | 0.22 |
| | Positive | 18 (40.9%) | 24 (51.1%) | 9 (75.0%) | | |
| 17R-RvD3 | Negative | 26 (59.1%) | 33 (70.2%) | 11 (91.7%) | 0.034** | 0.204 |
| | Positive | 18 (40.9%) | 14 (29.8%) | 1 (8.33%) | | |
| PD1 | | 0.30 [0.00;0.40] | 0.30 [0.00;0.50] | 0.10 [0.00;0.23] | 0.959 | 0.993 |
| 10S,17S-diHDHA | | 0.10 [0.00;0.52] | 0.40 [0.10;0.70] | 0.10 [0.00;0.52] | 0.197 | 0.425 |
| 17R-PD1 | Negative | 41 (93.2%) | 42 (89.4%) | 10 (83.3%) | 0.297 | 0.534 |
| | Positive | 3 (6.82%) | 5 (10.6%) | 2 (16.7%) | | |
| 22-OH-PD1 | Negative | 42 (95.5%) | 42 (89.4%) | 5 (41.7%) | <0.001** | 0.001 |
| | Positive | 2 (4.55%) | 5 (10.6%) | 7 (58.3%) | | |
| PCTR1 | Negative | 39 (88.6%) | 41 (87.2%) | 11 (91.7%) | 0.901 | 0.993 |
| | Positive | 5 (11.4%) | 6 (12.8%) | 1 (8.33%) | | |
| PCTR2 | Negative | 40 (90.9%) | 45 (95.7%) | 12 (100%) | 0.181 | 0.408 |
| | Positive | 4 (9.09%) | 2 (4.26%) | 0 (0.00%) | | |
| PCTR3 | Negative | 34 (77.3%) | 38 (80.9%) | 7 (58.3%) | 0.394 | 0.59 |
| | Positive | 10 (22.7%) | 9 (19.1%) | 5 (41.7%) | | |
| MaR1 | Negative | 34 (77.3%) | 24 (51.1%) | 5 (41.7%) | 0.005** | 0.05 |
| | Positive | 10 (22.7%) | 23 (48.9%) | 7 (58.3%) | | |
| 7S-14S-diHDHA | Negative | 27 (61.4%) | 34 (72.3%) | 7 (58.3%) | 0.727 | 0.921 |
| | Positive | 17 (38.6%) | 13 (27.7%) | 5 (41.7%) | | |
| MaR2 | | 2.70 [0.10;12.8] | 1.30 [0.15;15.0] | 0.45 [0.18;0.88] | 0.255 | 0.492 |
| 4S,14S-diHDHA | Negative | 22 (50.0%) | 32 (68.1%) | 8 (66.7%) | 0.115 | 0.356 |
| | Positive | 22 (50.0%) | 15 (31.9%) | 4 (33.3%) | | |
| 22-OH-MaR1 | Negative | 30 (68.2%) | 31 (66.0%) | 5 (41.7%) | 0.169 | 0.398 |

| | | | | | | |
|---|---|---|---|---|---|---|
| | Positive | 14 (31.8%) | 16 (34.0%) | 7 (58.3%) | | |
| MCTR1 | Negative | 42 (95.5%) | 43 (91.5%) | 11 (91.7%) | 0.494 | 0.667 |
| | Positive | 2 (4.55%) | 4 (8.51%) | 1 (8.33%) | | |
| MCTR2 | Negative | 44 (100%) | 45 (95.7%) | 11 (91.7%) | 0.092 | 0.311 |
| | Positive | 0 (0.00%) | 2 (4.26%) | 1 (8.33%) | | |
| MCTR3 | Negative | 38 (86.4%) | 40 (85.1%) | 11 (91.7%) | 0.781 | 0.925 |
| | Positive | 6 (13.6%) | 7 (14.9%) | 1 (8.33%) | | |
| **n-3 DPA metabolome** | | | | | | |
| RvT1 | | 0.15 [0.10;0.23] | 0.10 [0.00;0.25] | 0.10 [0.10;0.20] | 0.329 | 0.568 |
| RvT2 | | 0.70 [0.20;1.20] | 0.30 [0.10;0.90] | 0.10 [0.00;0.50] | 0.015** | 0.136 |
| RvT3 | Negative | 26 (59.1%) | 23 (48.9%) | 9 (75.0%) | 0.763 | 0.925 |
| | Positive | 18 (40.9%) | 24 (51.1%) | 3 (25.0%) | | |
| RvT4 | | 0.20 [0.00;0.40] | 0.10 [0.00;0.30] | 0.00 [0.00;0.10] | 0.04** | 0.217 |
| $RvD1_{n\text{-}3\ DPA}$ | | 0.20 [0.10;0.30] | 0.20 [0.10;0.30] | 0.10 [0.00;0.23] | 0.337 | 0.568 |
| $RvD2_{n\text{-}3\ DPA}$ | | 0.80 [0.48;1.10] | 0.30 [0.20;0.60] | 0.25 [0.08;0.43] | <0.001** | <0.001** |
| $RvD5_{n\text{-}3\ DPA}$ | Negative | 30 (68.2%) | 28 (59.6%) | 11 (91.7%) | 0.447 | 0.619 |
| | Positive | 14 (31.8%) | 19 (40.4%) | 1 (8.33%) | | |
| $PD1_{n\text{-}3\ DPA}$ | | 5.60 [0.00;13.6] | 0.30 [0.00;7.25] | 0.35 [0.00;2.60] | 0.156 | 0.398 |
| 10S,17S-diHDPA | Negative | 28 (63.6%) | 35 (74.5%) | 6 (50.0%) | 0.861 | 0.968 |
| | Positive | 16 (36.4%) | 12 (25.5%) | 6 (50.0%) | | |
| $MaR1_{n\text{-}3\ DPA}$ | Negative | 27 (61.4%) | 31 (66.0%) | 11 (91.7%) | 0.09 | 0.311 |
| | Positive | 17 (38.6%) | 16 (34.0%) | 1 (8.33%) | | |
| 7S,14S-diHDPA | | 0.80 [0.00;1.25] | 0.70 [0.00;1.30] | 0.05 [0.00;0.48] | 0.374 | 0.577 |
| **EPA metabolome** | | | | | | |
| RvE1 | | 0.85 [0.38;1.60] | 0.50 [0.05;1.60] | 0.30 [0.08;1.12] | 0.091 | 0.311 |
| RvE2 | | 4.90 [3.75;6.43] | 5.50 [3.80;7.90] | 4.70 [2.62;6.30] | 0.975 | 0.993 |
| RvE3 | Negative | 29 (65.9%) | 30 (63.8%) | 10 (83.3%) | 0.447 | 0.619 |
| | Positive | 15 (34.1%) | 17 (36.2%) | 2 (16.7%) | | |
| **AA metabolome** | | | | | | |
| $LXA_4$ | | 0.10 [0.00;0.10] | 0.10 [0.00;0.10] | 0.10 [0.00;0.10] | 0.533 | 0.702 |
| $LXB_4$ | | 3.80 [1.85;14.5] | 5.20 [1.45;36.2] | 6.60 [2.95;21.2] | 0.404 | 0.59 |
| 5S,15S-diHETE | | 6.10 [1.05;10.8] | 4.40 [1.90;7.90] | 6.35 [3.93;12.4] | 0.967 | 0.993 |
| 15-epi-$LXA_4$ | | 0.50 [0.30;0.70] | 0.20 [0.10;0.40] | 0.40 [0.18;0.52] | 0.005** | 0.05 |
| 15-epi-$LXB_4$ | | 77.2 [58.4;88.7] | 16.4 [0.25;36.6] | 0.30 [0.08;15.6] | <0.001** | <0.001** |
| $LTB_4$ | | 3.60 [1.37;12.2] | 6.50 [2.45;16.7] | 2.05 [0.40;5.50] | 0.805 | 0.925 |
| 5S, 12S-diHETE | Negative | 39 (88.6%) | 38 (80.9%) | 8 (66.7%) | 0.076 | 0.311 |

| | | | | | | |
|---|---|---|---|---|---|---|
| | Positive | 5 (11.4%) | 9 (19.1%) | 4 (33.3%) | | |
| 12-epi-$LTB_4$ | | 1.05 [0.50;3.32] | 1.60 [0.45;3.00] | 0.30 [0.27;1.20] | 0.363 | 0.577 |
| 6-trans, 12-epi-$LTB_4$ | | 1.10 [0.50;3.35] | 1.50 [0.50;3.00] | 0.30 [0.20;1.50] | 0.277 | 0.516 |
| 20-OH-$LTB_4$ | | 1.00 [0.18;5.65] | 2.70 [0.35;9.25] | 3.10 [0.70;9.18] | 0.162 | 0.398 |
| $LTC_4$ | Negative | 27 (61.4%) | 22 (46.8%) | 5 (41.7%) | 0.125 | 0.356 |
| | Positive | 17 (38.6%) | 25 (53.2%) | 7 (58.3%) | | |
| $LTD_4$ | | 0.80 [0.30;2.82] | 0.80 [0.00;4.95] | 0.60 [0.00;3.20] | 0.733 | 0.921 |
| $LTE_4$ | | 65.3 [38.3;188] | 156 [47.6;456] | 185 [75.4;1943] | 0.018** | 0.139 |
| $PGD_2$ | | 3.15 [0.55;4.23] | 2.40 [0.90;6.45] | 4.55 [2.08;10.3] | 0.166 | 0.398 |
| $PGE_2$ | | 35.1 [9.55;108] | 61.6 [28.1;207] | 72.0 [21.5;162] | 0.051 | 0.228 |
| $PGF_2$□ | | 72.2 [37.7;157] | 137 [67.0;387] | 179 [55.2;253] | 0.032** | 0.204 |
| $TxB_2$ | | 46.5 [12.6;70.3] | 54.6 [23.5;93.6] | 60.0 [12.1;110] | 0.215 | 0.446 |

All summary statistics are absolute counts (%) for categorical variables and median [inter-quartile range] for continuous data. n refers to the number of patients with non-missing data for the corresponding variable.

*Negative means lipid measurement under the detection limit. Positive means lipid measurement above the detection limit.

** Note that p value <0.05

**Supplementary Table 3: Summaries of lipid mediators' selection in the Lasso regression model for the disease severity**

| Lipid Mediators | Frequency* | OR (2 vs 1) | 95%CI (2 vs 1) | OR (3 vs 1) | 95%CI (3 vs 1) |
|---|---|---|---|---|---|
| 15-epi-$LXB_4$ | 1000 | 0.70 | [0.61 ;0.81] | 0.63 | [0.49 ;0.82] |
| $LXB_4$ | 1000 | 1.02 | [0.99 ;1.03] | 1.00 | [0.98 ;1.03] |
| $PGE_2$ | 1000 | 1.05 | [1.01 ;1.09] | 1.02 | [0.97 ;1.08] |
| 22-OH-PD1 (Neg/Pos) | 1000 | 2.50 | [0.46 ;13.61] | 29.40 | [4.74 ;182.3] |
| PD1 | 935 | 2.55 | [0.68 ;9.60] | 0.24 | [0.02 ;3.70] |
| $RvD5_{n\text{-}3\ DPA}$ (Neg/Pos) | 935 | 1.45 | [0.61 ;3.44] | 0.19 | [0.02 ;1.66] |
| $RvD2_{n\text{-}3\ DPA}$ | 830 | 0.15 | [0.05 ;0.50] | 0.17 | [0.03 ;1.02] |
| RvE2 | 830 | 1.05 | [0.92 ;1.20] | 0.84 | [0.64 ;1.13] |
| 15-epi$LXA_4$ | 683 | 0.07 | [0.01 ;0.39] | 0.49 | [0.09 ;2.74] |
| $LTD_4$ | 683 | 1.06 | [0.98 ;1.15] | 0.99 | [0.86 ;1.15] |
| $RvD1_{n\text{-}3\ DPA}$ | 683 | 1.76 | [0.50 ;6.18] | 0.23 | [0.01 ;8.85] |
| 20-OH-$LTB_4$ | 519 | 1.00 | [0.97 ;1.04] | 1.02 | [0.99 ;1.06] |
| MaR1 (Neg/Pos) | 519 | 3.26 | [1.31 ;8.08] | 4.76 | [1.24 ;18.30] |
| RvD2 (Neg/Pos) | 519 | 0.67 | [0.28 ;1.61] | 1.13 | [0.31 ;4.16] |
| RvT3 (Neg/Pos) | 519 | 1.50 | [0.66 ;3.46] | 0.48 | [0.11 ;2.03] |
| 10S,17S-diHDPA (Neg/Pos) | 369 | 1.50 | [0.75 ;2.98] | 0.84 | [0.27 ;3.31] |
| 7S,14S-diHDHA (Neg/Pos) | 206 | 0.61 | [0.25 ;1.47] | 1.13 | [0.31 ;4.16] |
| RvE3 (Neg/Pos) | 206 | 1.10 | [0.46 ;2.59] | 0.39 | [0.07 ;2.00] |
| $LTE_4$ | 105 | 1.01 | [1.00 ;1.02] | 1.01 | [1.00 ;1.02] |
| 17R-RvD3 (Neg/Pos) | 56 | 0.61 | [0.26 ;1.46] | 0.13 | [0.02 ;1.11] |

* High frequency represents the likely chosen as the strong predictors of the disease severity.
CI, confidence interval; OR, odds ratio

### Supplemental Table 4: Summary Table Survivors/Non-survivors

| Characteristic | n | Summary statistic<br>Survivor (N=95) | n | Summary statistic<br>Non-survivor (N=8) |
|---|---|---|---|---|
| Gender – no. (%) | 95 | | 7 | |
| - Male | | 62/95 (65%) | | 2/7 (29%) |
| - Female | | 33/95 (35%) | | 5/7 (71%) |
| Age (years) – median (IQR) | 95 | 40.00(30.00,49.00) | 7 | 41.00(38.50,49.00) |
| Weight (kg) – median (IQR) | 95 | 50.00(45.25,60.00) | 7 | 58.50(48.25,60.00) |
| Glasgow coma score – median (IQR) | 95 | 15.00(13.00,15.00) | 8 | 11.50(8.00,14.00) |
| *Diagnostic category – no. (%) | 95 | | 8 | |
| - definite TBM | | 76/95 (80%) | | 5/8 (62%) |
| - possible TBM | | 14/95 (15%) | | 1/8 (12%) |
| - probable TBM | | 5/95 (5%) | | 2/8 (25%) |
| Treatment arm – no. (%) | 95 | | 8 | |
| - Aspirin_1000mg | | 32/95 (34%) | | 0/8 (0%) |
| - Aspirin_81mg | | 30/95 (32%) | | 5/8 (62%) |
| - Placebo | | 33/95 (35%) | | 3/8 (38%) |

All summary statistics are absolute counts (%) for categorical variables and median (inter-quartile range = IQR) for continuous data. n refers to the number of patients with non-missing data for the corresponding variable.

### Supplemental Table 5: Comparison of lipid mediators by survival outcome

| Lipid Mediators | | Survivors (N=95) | Non-Survivor (N=8) | P Overall | Adjusted P |
|---|---|---|---|---|---|
| **DHA metabolome** | | | | | |
| RvD1 | Negative* | 59 (62.1%) | 5 (62.5%) | 1 | 1 |
| | Positive | 36 (37.9%) | 3 (37.5%) | | |
| RvD2 | Negative | 62 (65.3%) | 5 (62.5%) | 1 | 1 |
| | Positive | 33 (34.7%) | 3 (37.5%) | | |
| RvD3 | Negative | 69 (72.6%) | 5 (62.5%) | 0.684 | 0.939 |
| | Positive | 26 (27.4%) | 3 (37.5%) | | |
| RvD4 | | 0.30 [0.10;0.50] | 0.15 [0.00;0.20] | 0.077 | 0.258 |
| RvD5 | | 0.60 [0.20;1.15] | 0.10 [0.00;0.23] | 0.028** | 0.179 |
| RvD6 | Negative | 62 (65.3%) | 6 (75.0%) | 0.374 | 0.939 |
| | Positive | 33 (34.7%) | 2 (25.0%) | | |
| 17R-RvD1 | Negative | 49 (51.6%) | 3 (37.5%) | 0.488 | 0.826 |
| | Positive | 46 (48.4%) | 5 (62.5%) | | |
| 17R-RvD3 | Negative | 62 (65.3%) | 8 (100%) | 0.052 | 0.202 |
| | Positive | 33 (34.7%) | 0 (0.00%) | | |
| PD1 | | 0.30 [0.00;0.50] | 0.00 [0.00;0.08] | 0.013** | 0.114 |
| 10S,17S-diHDHA | | 0.30 [0.00;0.60] | 0.10 [0.00;0.43] | 0.374 | 0.784 |
| 17R-PD1 | Negative | 86 (90.5%) | 7 (87.5%) | 0.572 | 0.867 |
| | Positive | 9 (9.47%) | 1 (12.5%) | | |
| 22-OH-PD1 | Negative | 82 (86.3%) | 7 (87.5%) | 1 | 1 |
| | Positive | 13 (13.7%) | 1 (12.5%) | | |
| PCTR1 | Negative | 83 (87.4%) | 8 (100%) | 0.591 | 0.867 |
| | Positive | 12 (12.6%) | 0 (0.00%) | | |
| PCTR2 | Negative | 74 (77.9%) | 5 (62.5%) | 0.392 | 0.784 |
| | Positive | 21 (22.1%) | 3 (37.5%) | | |
| PCTR3 | Negative | 34 (77.3%) | 38 (80.9%) | 0.385 | 0.784 |
| | Positive | 10 (22.7%) | 9 (19.1%) | | |
| MaR1 | Negative | 58 (61.1%) | 5 (62.5%) | 1 | 1 |
| | Positive | 37 (38.9%) | 3 (37.5%) | | |
| 7S,14S-diHDHA | Negative | 27 (61.4%) | 34 (72.3%) | 0.727 | 0.939 |
| | Positive | 17 (38.6%) | 13 (27.7%) | | |

| | | | | | |
|---|---|---|---|---|---|
| MaR2 | | 2.20 [0.20;13.8] | 0.05 [0.00;0.20] | 0.002** | 0.05 |
| 4S,14S-diHDHA | Negative | 56 (58.9%) | 6 (75.0%) | 0.742 | 0.826 |
| | Positive | 39 (41.1%) | 2 (25.0%) | | |
| 22-OH-MaR1 | Negative | 61 (64.2%) | 5 (62.5%) | 1 | 1 |
| | Positive | 34 (35.8%) | 3 (37.5%) | | |
| MCTR1 | Negative | 88 (92.6%) | 8 (100%) | 1 | 1 |
| | Positive | 7 (7.37%) | 0 (0.00%) | | |
| MCTR2 | Negative | 92 (96.8%) | 8 (100%) | 1 | 1 |
| | Positive | 3 (3.16%) | 0 (0.00%) | | |
| MCTR3 | Negative | 81 (85.3%) | 8 (100%) | 0.594 | 0.867 |
| | Positive | 14 (14.7%) | 0 (0.00%) | | |
| **n-3 DPA metabolome** | | | | | |
| RvT1 | | 0.10 [0.05;0.25] | 0.15 [0.10;0.20] | 0.635 | 0.902 |
| RvT2 | | 0.40 [0.15;1.20] | 0.05 [0.00;0.10] | 0.003** | 0.05 |
| RvT3 | Negative | 52 (54.7%) | 6 (75.0%) | 0.461 | 0.826 |
| | Positive | 43 (45.3%) | 2 (25.0%) | | |
| RvT4 | | 0.10 [0.00;0.30] | 0.00 [0.00;0.10] | 0.05** | 0.202 |
| $RvD1_{n-3\ DPA}$ | | 0.20 [0.10;0.30] | 0.05 [0.00;0.25] | 0.256 | 0.602 |
| $RvD2_{n-3\ DPA}$ | | 0.50 [0.30;0.80] | 0.20 [0.00;0.32] | 0.008** | 0.09 |
| $RvD5_{n-3\ DPA}$ | Negative | 61 (64.2%) | 8 (100%) | 0.05** | 0.202 |
| | Positive | 34 (35.8%) | 0 (0.00%) | | |
| $PD1_{n-3\ DPA}$ | | 0.60 [0.00;9.60] | 0.15 [0.00;0.32] | 0.086 | 0.273 |
| 10S,17S-diHDPA | Negative | 63 (66.3%) | 6 (75.0%) | 1 | 1 |
| | Positive | 32 (33.7%) | 2 (25.0%) | | |
| $MaR1_{n-3\ DPA}$ | Negative | 64 (67.4%) | 5 (62.5%) | 1 | 1 |
| | Positive | 31 (32.6%) | 3 (37.5%) | | |
| 7S,14S-diHDPA | | 0.70 [0.00;1.30] | 0.00 [0.00;0.05] | 0.053 | 0.202 |
| **EPA metabolome** | | | | | |
| RvE1 | | 0.70 [0.20;1.60] | 0.15 [0.00;1.02] | 0.127 | 0.361 |
| RvE2 | | 5.10 [3.65;6.50] | 5.85 [3.62;6.52] | 0.897 | 1 |
| RvE3 | Negative | 63 (66.3%) | 6 (75.0%) | 1 | 1 |
| | Positive | 32 (33.7%) | 2 (25.0%) | | |
| **AA metabolome** | | | | | |
| $LXA_4$ | | 0.10 [0.00;0.10] | 0.05 [0.00;0.10] | 0.161 | 0.4334 |

| | | | | | |
|---|---|---|---|---|---|
| $LXB_4$ | | 4.20 [1.80;28.1] | 4.65 [2.72;6.45] | 0.49 | 0.826 |
| 5S,15S-diHETE | | 5.40 [1.90;10.2] | 7.75 [4.67;8.23] | 0.591 | 0.867 |
| 15-epi-$LXA_4$ | | 0.30 [0.20;0.60] | 0.45 [0.35;0.60] | 0.515 | 0.842 |
| 15-epi$LXB_4$ | | 37.5 [10.2;77.3] | 0.00 [0.00;0.20] | <0.001** | 0.02** |
| $LTB_4$ | | 5.00 [1.65;16.7] | 2.10 [0.92;3.65] | 0.07 | 0.251 |
| 5S,12S-diHETE | Negative | 81 (85.3%) | 4 (50.0%) | 0.03** | 0.179 |
| | Positive | 14 (14.7%) | 4 (50.0%) | | |
| 12-epi-$LTB_4$ | | 1.30 [0.45;3.10] | 0.35 [0.20;0.60] | 0.037** | 0.2 |
| 6-trans,12-epi-$LTB_4$ | | 1.10 [0.45;3.10] | 0.35 [0.30;1.30] | 0.092 | 0.276 |
| 20-OH-$LTB_4$ | | 1.40 [0.25;8.60] | 1.75 [0.75;5.52] | 0.753 | 0.968 |
| $LTC_4$ | Negative | 51 (53.7%) | 3 (37.5%) | 0.473 | 0.826 |
| | Positive | 44 (46.3%) | 5 (62.5%) | | |
| $LTD_4$ | | 0.80 [0.20;3.05] | 1.00 [0.00;11.9] | 0.98 | 1 |
| $LTE_4$ | | 92.7 [43.3;335] | 827 [524;1305] | 0.007** | 0.09 |
| $PGD_2$ | | 2.70 [0.90;4.35] | 7.10 [5.27;14.0] | 0.015 | 0.116 |
| $PGE_2$ | | 45.7 [17.5;168] | 33.9 [3.38;61.7] | 0.198 | 0.508 |
| PGF[illegible] | | 111 [48.4;261] | 84.6 [22.3;141] | 0.229 | 0.563 |
| $TXB_2$ | | 48.7 [16.2;85.0] | 78.6 [49.9;104] | 0.358 | 0.784 |

All summary statistics are absolute counts (%) for categorical variables and median [inter-quartile range] for continuous data. n refers to the number of patients with non-missing data for the corresponding variable.

*Negative means lipid measurement under the detection limit. Positive means lipid measurement above the detection limit.

** Note that p value <0.05

**Supplementary Table 6: Summaries of lipid mediators' selection in the Lasso regression model for survival outcome**

| Lipid Mediators | Frequency | OR | 95% CI (OR) |
|---|---|---|---|
| $PGD_2$ | 998 | 1.26 | [1.10; 1.52] |
| 15-epi-$LXB_4$ | 984 | 0.25 | [0.02; 0.67] |
| 17R-RvD3 (Neg/Pos) | 974 | 0.00 | NA |
| 5S,12S-diHETE (Neg/Pos) | 953 | 5.78 | [1.24; 27.20] |
| $RvD5_{n\text{-}3\ DPA}$ (Neg/Pos) | 938 | 0.00 | NA |
| PD1 | 904 | 0.01 | [0.00; 0.37] |
| PCTR2 (Neg/Pos) | 860 | 2.57 | [0.13; 19.22] |
| RvT3 (Neg/Pos) | 785 | 0.40 | [0.06; 1.85] |
| PCTR1 (Neg/Pos) | 695 | 0.00 | NA |
| MCTR1 (Neg/Pos) | 610 | 0.00 | NA |
| RvD6 (Neg/Pos) | 610 | 0.63 | [0.09; 2.89] |
| $PGF_{2}$□ | 528 | 0.96 | [0.87; 1.00] |
| $RvD1_{n\text{-}3\ DPA}$ | 456 | 0.210 | [0.00; 2.84] |
| 10S,17S-diHDPA (Neg/Pos) | 378 | 0.66 | [0.09; 3.04] |
| $LTC_4$ (Neg/Pos) | 302 | 1.93 | [0.45; 9.85] |
| $LXB_4$ | 165 | 0.94 | [0.81; 1.00] |
| MaR1 (Neg/Pos) | 165 | 0.94 | [0.18; 4.07] |
| $PD1_{n\text{-}3\ DPA}$ | 165 | 0.57 | [0.00; 0.90] |
| RvT2 | 115 | 0.02 | [0.00; 0.43] |

### Supplemental Table 7: Summary table placebo, 81 mg and 1000mg aspirin

| Characteristic | n | Summary statistic Placebo (N=36) | n | Summary statistic 81mg aspirin (N=35) | n | Summary statistic 1000mg aspirin (N=32) |
|---|---|---|---|---|---|---|
| Gender – no. (%) | 36 | | 34 | | 32 | |
| - Male | | 24/36 (67%) | | 24/34 (71%) | | 16/32 (50%) |
| - Female | | 12/36 (33%) | | 10/34 (29%) | | 16/32 (50%) |
| Age (years) – median (IQR) | 36 | 42.00(32.75,50.00) | 34 | 39.00(32.50,47.75) | 32 | 39.50(29.75,51.25) |
| Weight (kg) – median (IQR) | 36 | 50.75(44.75,60.00) | 34 | 50.50(45.00,58.88) | 32 | 50.00(46.38,60.50) |
| Glasgow coma score – median (IQR) | 36 | 14.00(12.75,15.00) | 35 | 14.00(12.50,15.00) | 32 | 15.00(12.75,15.00) |
| Diagnostic category – no. (%) | 36 | | 35 | | 32 | |
| - definite TBM | | 31/36 (86%) | | 26/35 (74%) | | 24/32 (75%) |
| - possible TBM | | 3/36 (8%) | | 5/35 (14%) | | 7/32 (22%) |
| - probable TBM | | 2/36 (6%) | | 4/35 (11%) | | 1/32 (3%) |

All summary statistics are absolute counts (%) for categorical variables and median (inter-quartile range = IQR) for continuous data. n refers to the number of patients with non-missing data for the corresponding variable.

**Supplemental Table 8**: **Comparison of lipid mediator reduction after 30 days between placebo and 81mg aspirin**

| Lipid Mediators | Placebo (N=34) | Aspirin 81mg (N=28) | P overall | Adjusted P |
|---|---|---|---|---|
| **DHA metabolome** | | | | |
| RvD1 | 0.10 [0.00;0.20] | 0.10 [0.00;0.20] | 0.842 | 0.965 |
| RvD2 | 1.10 [0.23;2.10] | 1.20 [0.38;2.97] | 0.332 | 0.965 |
| RvD3 | 0.00 [0.00;0.10] | 0.00 [0.00;0.00] | 0.306 | 0.965 |
| RvD4 | 0.10 [0.00;0.20] | 0.10 [0.00;0.20] | 0.7 | 0.965 |
| RvD5 | 0.10 [0.00;0.10] | 0.10 [0.00;0.20] | 0.665 | 0.965 |
| RvD6 | 0.00 [0.00;0.10] | 0.00 [0.00;0.10] | 0.822 | 0.965 |
| 17R-RvD1 | 0.00 [0.00;0.10] | 0.00 [0.00;0.10] | 0.538 | 0.965 |
| 17R-RvD3 | 0.00 [0.00;0.00] | 0.00 [0.00;0.00] | 0.773 | 0.965 |
| PD1 | 0.20 [0.02;0.30] | 0.20 [0.18;0.30] | 0.857 | 0.965 |
| 10S,17S-diHDHA | 0.00 [0.00;0.20] | 0.00 [0.00;0.10] | 0.446 | 0.965 |
| 17R-PD1 | 0.00 [0.00;0.10] | 0.00 [0.00;0.00] | 0.207 | 0.965 |
| 22-OH-PD1 | 0.40 [0.10;0.60] | 0.35 [0.18;0.62] | 0.892 | 0.965 |
| PCTR1 | 0.00 [0.00;0.00] | 0.00 [0.00;0.00] | 0.842 | 0.965 |
| PCTR2 | 0.00 [0.00;0.00] | 0.00 [0.00;0.00] | 0.364 | 0.965 |
| PCTR3 | 0.00 [0.00;0.00] | 0.00 [0.00;0.00] | 0.975 | 1 |
| MaR1 | 0.10 [0.00;0.18] | 0.05 [0.00;0.10] | 0.173 | 0.965 |
| 7S,14S-diHDHA | 0.00 [0.00;0.15] | 0.00 [0.00;0.42] | 0.543 | 0.965 |
| MaR2 | 1.00 [0.20;12.4] | 1.00 [0.10;8.30] | 0.702 | 0.965 |
| 4S,14S-diHDHA | 0.00 [0.00;0.00] | 0.00 [0.00;0.02] | 1 | 1 |
| 22-OH-MaR1 | 0.35 [0.00;0.80] | 0.40 [0.00;0.70] | 0.885 | 0.965 |
| MCTR1 | | | | |
| MCTR2 | 0.00 [0.00;0.00] | 0.00 [0.00;0.00] | 0.364 | 0.965 |
| MCTR3 | 0.00 [0.00;0.00] | 0.00 [0.00;0.00] | 0.945 | 1 |
| **n-3 DPA metabolome** | | | | |
| RvT1 | 0.10 [0.00;0.10] | 0.10 [0.00;0.10] | 0.754 | 0.965 |
| RvT2 | 0.30 [0.10;1.35] | 0.15 [0.10;1.12] | 0.538 | 0.965 |
| RvT3 | 0.00 [0.00;0.00] | 0.00 [0.00;0.10] | 0.297 | 0.965 |
| RvT4 | 0.00 [0.00;0.10] | 0.05 [0.00;0.10] | 0.875 | 0.965 |
| $RvD1_{n\text{-}3\ DPA}$ | 0.00 [0.00;0.10] | 0.10 [0.00;0.10] | 0.763 | 0.965 |
| $RvD2_{n\text{-}3\ DPA}$ | 0.10 [0.00;0.10] | 0.00 [0.00;0.10] | 0.452 | 0.965 |
| $RvD5_{n\text{-}3\ DPA}$ | 0.00 [0.00;0.08] | 0.00 [0.00;0.20] | 0.16 | 0.965 |

| | | | | |
|---|---|---|---|---|
| $PD1_{n\text{-}3\ DPA}$ | 4.00 [1.33;10.1] | 6.20 [0.08;8.48] | 0.994 | 1 |
| 10S,17S-diHDPA | 5.95 [0.00;11.8] | 3.25 [0.00;6.03] | 0.156 | 0.965 |
| $MaR1_{n\text{-}3\ DPA}$ | 0.00 [0.00;0.15] | 0.20 [0.00;0.52] | 0.036** | 0.965 |
| 7S,14S-diHDPA | 0.30 [0.00;1.58] | 0.65 [0.10;2.05] | 0.177 | 0.965 |
| **EPA metabolome** | | | | |
| RvE1 | 0.15 [0.10;0.30] | 0.20 [0.08;0.20] | 0.805 | 0.965 |
| RvE2 | 2.70 [1.92;3.98] | 2.85 [2.00;3.80] | 0.882 | 0.965 |
| RvE3 | 0.00 [0.00;0.20] | 0.00 [0.00;0.05] | 0.577 | 0.965 |
| **AA metabolome** | | | | |
| $LXA_4$ | 0.00 [0.00;0.10] | 0.00 [0.00;0.02] | 0.655 | 0.965 |
| $LXB_4$ | 1.75 [1.30;5.70] | 2.55 [1.82;5.35] | 0.186 | 0.965 |
| 5S,15S-diHETE | 3.30 [2.35;5.90] | 3.55 [1.60;4.62] | 0.547 | 0.965 |
| 15-epi-$LXA_4$ | 0.10 [0.00;0.20] | 0.10 [0.00;0.10] | 0.192 | 0.965 |
| 15-epi-$LXB_4$ | 12.1 [7.02;23.5] | 16.6 [7.40;31.1] | 0.687 | 0.965 |
| $LTB_4$ | 0.70 [0.20;2.57] | 0.50 [0.30;1.35] | 0.837 | 0.965 |
| 5S,12S-diHETE | 0.00 [0.00;0.00] | 0.00 [0.00;0.00] | 0.541 | 0.965 |
| 12-epi-$LTB_4$ | 0.15 [0.00;0.50] | 0.10 [0.00;0.30] | 0.667 | 0.965 |
| 6-trans,12-epi-$LTB_4$ | 0.10 [0.02;0.30] | 0.15 [0.08;0.40] | 0.503 | 0.965 |
| 20-OH-$LTB_4$ | 12.1 [7.02;23.5] | 16.6 [7.40;31.1] | 0.666 | 0.965 |
| $LTC_4$ | 0.00 [0.00;0.27] | 0.00 [0.00;0.12] | 0.717 | 0.965 |
| $LTD_4$ | 0.70 [0.02;1.10] | 0.70 [0.15;2.30] | 0.462 | 0.965 |
| $LTE_4$ | 9.15 [2.68;28.6] | 7.45 [2.85;100] | 0.815 | 0.965 |
| $PGD_2$ | 3.35 [1.72;4.97] | 2.10 [1.50;3.35] | 0.099 | 0.965 |
| $PGE_2$ | 4.30 [1.65;22.0] | 3.70 [1.28;18.7] | 0.756 | 0.965 |
| $PGF_{2}$□ | 11.8 [7.05;30.1] | 12.7 [5.62;26.6] | 0.729 | 0.965 |
| $TxB_2$ | 16.2 [6.97;30.6] | 8.20 [3.53;19.7] | 0.058 | 0.965 |

** Note that p value <0.05

**Supplemental Table 9**: **Comparison of lipid mediator reduction after 30 days between placebo and 1000mg aspirin**

| Lipid Mediators | Placebo (N=34) | 1000mg aspirin (N=27) | P overall | Adjusted P |
|---|---|---|---|---|
| **DHA metabolome** | | | | |
| RvD1 | 0.10 [0.00;0.20] | 0.10 [0.00;0.20] | 0.258 | 0.648 |
| RvD2 | 1.10 [0.23;2.10] | 1.80 [0.40;2.95] | 0.292 | 0.656 |
| RvD3 | 0.00 [0.00;0.10] | 0.00 [0.00;0.10] | 0.186 | 0.648 |
| RvD4 | 0.10 [0.00;0.20] | 0.10 [0.00;0.20] | 0.897 | 0.914 |
| RvD5 | 0.10 [0.00;0.10] | 0.10 [0.00;0.25] | 0.238 | 0.648 |
| RvD6 | 0.00 [0.00;0.10] | 0.00 [0.00;0.10] | 0.781 | 0.914 |
| 17R-RvD1 | 0.00 [0.00;0.10] | 0.00 [0.00;0.00] | 0.278 | 0.654 |
| 17R-RvD3 | 0.00 [0.00;0.00] | 0.00 [0.00;0.05] | 0.437 | 0.786 |
| PD1 | 0.20 [0.02;0.30] | 0.20 [0.10;0.40] | 0.637 | 0.905 |
| 10S,17S-diHDHA | 0.00 [0.00;0.20] | 0.10 [0.00;0.20] | 0.433 | 0.786 |
| 17R-PD1 | 0.00 [0.00;0.10] | 0.00 [0.00;0.00] | 0.584 | 0.866 |
| 22-OH-PD1 | 0.40 [0.10;0.60] | 0.40 [0.35;0.60] | 0.308 | 0.666 |
| PCTR1 | 0.00 [0.00;0.00] | 0.00 [0.00;0.00] | 0.238 | 0.648 |
| PCTR2 | 0.00 [0.00;0.00] | 0.00 [0.00;0.00] | 0.211 | 0.648 |
| PCTR3 | 0.00 [0.00;0.00] | 0.00 [0.00;0.00] | 0.691 | 0.914 |
| MaR1 | 0.10 [0.00;0.18] | 0.00 [0.00;0.15] | 0.093 | 0.648 |
| 7S,14S-diHDHA | 0.00 [0.00;0.15] | 0.00 [0.00;0.00] | 0.49 | 0.827 |
| MaR2 | 1.00 [0.20;12.4] | 1.00 [0.40;21.5] | 0.827 | 0.914 |
| 4S,14S-diHDHA | 0.00 [0.00;0.00] | 0.00 [0.00;0.05] | 0.885 | 0.914 |
| 22-OH-MaR1 | 0.35 [0.00;0.80] | 0.60 [0.25;0.85] | 0.264 | 0.648 |
| MCTR1 | 0.00 [0.00;0.00] | 0.00 [0.00;0.00] | 0.11 | 0.648 |
| MCTR2 | 0.00 [0.00;0.00] | 0.00 [0.00;0.00] | 0.373 | 0.767 |
| MCTR3 | 0.00 [0.00;0.00] | 0.00 [0.00;0.00] | 0.826 | 0.914 |
| **n-3 DPA metabolome** | | | | |
| RvT1 | 0.10 [0.00;0.10] | 0.10 [0.00;0.10] | 0.894 | 0.914 |
| RvT2 | 0.30 [0.10;1.35] | 0.20 [0.10;1.20] | 0.849 | 0.914 |
| RvT3 | 0.00 [0.00;0.00] | 0.00 [0.00;0.10] | 0.255 | 0.648 |
| RvT4 | 0.00 [0.00;0.10] | 0.10 [0.00;0.15] | 0.434 | 0.786 |
| $RvD1_{n\text{-}3\ DPA}$ | 0.00 [0.00;0.10] | 0.00 [0.00;0.10] | 0.885 | 0.914 |
| $RvD2_{n\text{-}3\ DPA}$ | 0.10 [0.00;0.10] | 0.00 [0.00;0.10] | 0.383 | 0.767 |
| $RvD5_{n\text{-}3\ DPA}$ | 0.00 [0.00;0.08] | 0.00 [0.00;0.40] | 0.121 | 0.648 |

| | | | | |
|---|---|---|---|---|
| $PD1_{n-3\ DPA}$ | 4.00 [1.33;10.1] | 4.80 [1.00;8.70] | 0.856 | 0.914 |
| 10S,17S-diHDPA | 5.95 [0.00;11.8] | 6.00 [1.65;10.9] | 0.49 | 0.827 |
| $MaR1_{n-3\ DPA}$ | 0.00 [0.00;0.15] | 0.00 [0.00;0.00] | 0.665 | 0.914 |
| 7S,14S-diHDPA | 0.30 [0.00;1.58] | 1.20 [0.20;2.10] | 0.125 | 0.648 |
| **EPA metabolome** | | | | |
| RvE1 | 0.15 [0.10;0.30] | 0.20 [0.10;0.20] | 0.847 | 0.914 |
| RvE2 | 2.70 [1.92;3.98] | 2.70 [2.10;4.80] | 0.591 | 0.866 |
| RvE3 | 0.00 [0.00;0.20] | 0.00 [0.00;0.00] | 0.24 | 0.648 |
| **AA metabolome** | | | | |
| $LXA_4$ | 0.00 [0.00;0.10] | 0.00 [0.00;0.10] | 0.594 | 0.866 |
| $LXB_4$ | 1.75 [1.30;5.70] | 3.40 [2.25;4.30] | 0.069 | 0.648 |
| 5S,15S-diHETE | 3.30 [2.35;5.90] | 3.20 [2.25;5.25] | 0.733 | 0.914 |
| 15-epi-$LXA_4$ | 0.10 [0.00;0.20] | 0.10 [0.00;0.25] | 0.539 | 0.856 |
| 15-epi-$LXB_4$ | 12.1 [7.02;23.5] | 21.0 [11.7;33.0] | 0.23 | 0.648 |
| $LTB_4$ | 0.70 [0.20;2.57] | 2.20 [0.70;9.70] | 0.02** | 0.279 |
| 5S,12S-diHETE | 0.00 [0.00;0.00] | 0.00 [0.00;0.00] | 0.937 | 0.937 |
| 12-epi-$LTB_4$ | 0.15 [0.00;0.50] | 0.40 [0.10;1.45] | 0.021** | 0.279 |
| 6-trans,12-epi-$LTB_4$ | 0.10 [0.02;0.30] | 0.40 [0.10;1.10] | 0.015** | 0.279 |
| 20-OH-$LTB_4$ | 12.1 [7.02;23.5] | 21.0 [11.7;33.0] | 0.234 | 0.648 |
| $LTC_4$ | 0.00 [0.00;0.27] | 0.10 [0.00;0.30] | 0.238 | 0.648 |
| $LTD_4$ | 0.70 [0.02;1.10] | 0.90 [0.35;1.95] | 0.139 | 0.648 |
| $LTE_4$ | 9.15 [2.68;28.6] | 10.1 [5.80;40.5] | 0.537 | 0.856 |
| $PGD_2$ | 3.35 [1.72;4.97] | 1.80 [1.10;4.00] | 0.095 | 0.648 |
| $PGE_2$ | 4.30 [1.65;22.0] | 5.40 [1.60;13.3] | 0.816 | 0.914 |
| $PGF_{2a}$ | 11.8 [7.05;30.1] | 11.6 [6.15;33.0] | 0.805 | 0.914 |
| $TxB_2$ | 16.2 [6.97;30.6] | 1.10 [0.50;2.00] | <0.001** | <0.001** |

** Note that p.value <0.05
